# AAV delivery of *GBA1* suppresses α-synuclein accumulation in Parkinson’s disease models and restores motor dysfunction in a Gaucher’s disease model

**DOI:** 10.1101/2024.06.25.599664

**Authors:** Takuro Okai, Sho Sato, Mugdha Deshpande, Shin-ichi Matsumoto, Miyu Nakayama, Syunsuke Yamamoto, Bettina Strack-Logue, Takeshi Hioki, Maiko Tanaka, Gabriele Proetzel

## Abstract

Biallelic mutations in the *glucosylceramidase beta 1* (*GBA1*) gene are the underlying genetic cause of Gaucher’s disease (GD), resulting in a deficient lysosomal hydrolase and subsequent accumulation of glycosphingolipids. More recently, *GBA1* mutations have been identified as the most prevalent genetic risk factor for Parkinson’s disease (PD), associated with more pronounced symptoms characterized by earlier onset and accelerated cognitive decline. In these GBA-associated PD patients the α-synuclein pathology is more prominent, and recent data suggest a link between α-synucleinopathies and *GBA1* mutations. Here, we explored the effect of *GBA1* gene supplementation on the GD phenotypes and α-synuclein pathology by using the adeno-associated virus (AAV) system. We have compared two AAV serotypes, AAV5 and AAV9, and two different ubiquitous promoters, and demonstrate that both promoters work efficiently albeit not the same *in vitro* and *in vivo. GBA1* overexpression reduces the accumulation of glucosylsphingosine (GlcSph) and restores motor dysfunction in a GD mouse model. We further demonstrate that *GBA1* overexpression can dissolve phospho-α-synuclein aggregation induced by the addition of α-synuclein pre-formed fibril (PFF) in a mouse primary neuron model suggesting the direct effect of β-Glucocerebrosidase (GCase) on α-synuclein accumulation. *In vivo*, we show that GCase inhibition can induce insoluble high-molecular-weight α-synuclein aggregation and that delivery of *GBA1* achieves robust reduction of the α-synuclein aggregates in the mouse brain. In summary, GCase expression not only reduces GlcSph, but also restores GD motor dysfunction and removes α-synuclein aggregates which are the hallmark for PD and α-synucleinopathies. AAV delivery of *GBA1* is a powerful approach to restore glucocerebrosidase function and to resolve misfolded α-synuclein protein, with applications for GD and PD.

## Introduction

Glucosylceramidase beta 1 *(GBA1)* encodes for the lysosomal enzyme glucocerebrosidase (GCase, acid β-glucosidase) that cleaves the beta-glycosidic linkage of glycolipid glucosylceramide (GlcCer) to ceramide and glucose. GCase is critical for maintaining glycosphingolipid homeostasis. Around 500 different mutations have been identified in the *GBA1* gene, which have different degrees of impact on GCase function, from reduced expression, protein instability, disruption of lysosomal targeting and/or reduced enzyme activity [1–3]. *GBA1* has been established as the causal gene for Gaucher’s disease (GD), the most common sphingolipidosis lysosomal storage disorder. GD is an autosomal recessive disorder caused by the partial or complete deficiency of GCase activity which interrupts the hydrolysis reaction leading to the accumulation of GlcCer and its deacylated lysolipid glucosylsphingosine (GlcSph) [4]. Based on the broad spectrum of phenotypes, GD has historically been classified into three clinical subtypes, Gaucher type 1, 2, and 3 (GD1, GD2, and GD3, respectively). GD1, also called non-neuronopathic GD, accounts for ∼94% of all cases in the US, Europe, and Israel [5, 6]. The main clinical manifestations of GD1 are hepatomegaly, splenomegaly, fatigue, thrombocytopenia, and bone and pulmonary involvement. GD2 and GD3 patients present with additional central nervous system (CNS) involvement including acute and subacute progressive neurological decline, resulting in significantly reduced life expectancy [7, 8]. The association between mutations in *GBA1* and the development of Parkinson’s disease (PD) was first described in the 1990s, establishing pathogenic *GBA1* mutations as the most important genetic risk factor for PD [9–11]. Approximately 5–20% of PD patients carry a heterozygous *GBA1* mutation (GBA-PD), differentiated by a 30–40% reduction in GCase activity, earlier disease onset including an increased risk for developing dementia with an up to five years earlier onset [12–23]. The reduced GCase activity can be detected in the cerebrospinal fluid (CSF) and blood of GBA-PD patients [24–26], as well as in post mortem brain tissues [27]. In GBA-PD, GCase activity was the primary variable determining GlcSph levels which in turn shows a strong correlation with α-synuclein pathology in all brain areas other than the cerebellum, a brain region that does not exhibit PD pathology [27]. *In vitro* and *in vivo* models have been developed based on expression of pathogenic *GBA1* mutants or administration of the GCase inhibitor conduritol β epoxide (CBE), to investigate the impact of reduced GCase levels on glycosphingolipid accumulation, lysosomal dysfunction, and increased α-synuclein aggregation [28, 29]. In α-synuclein aggregate rodent models of PD, overexpression of *GBA1* has been demonstrated to have a neuroprotective effect by reducing oligomeric and aggregated forms of α-synuclein, restoring autophagy and lysosomal pathways, as well as preventing α-synuclein-mediated degeneration of dopamine neurons [30–33]. Consequently, *GBA1* supplementation is thought as a disease-modifying therapeutic approach to restore lysosomal homeostasis and reduce α-synuclein neuropathology for the treatment of PD and other α-synuclein-associated neurodegenerative diseases, such as multiple system atrophy, dementia with Lewy body, and Alzheimer’s disease [2, 34].

One effective approach to deliver *GBA1* is to use adeno-associated virus (AAV) as a vehicle with the goal to achieve *GBA1* expression in relevant cell types to slow or stop disease progression of GD and GBA-PD [35–37]. There are several AAV serotypes with good neuronal transduction profile some of which have been used in clinical trials targeting CNS diseases, e.g., AAV2, AAV5, AAV9, and AAVrh10 [38–42]. As it has been established that the combination of AAV serotype, choice of promoter, and route of administration affect tissue tropism and cellular expression, the understanding of the interaction of these elements is instrumental to clinical benefit [43–46].

In this paper we compare two AAV serotypes, AAV5 and AAV9, and two different ubiquitous promoters, the chicken β-actin hybrid promoter (CBh) and the short form for the elongation factor 1α promoter (EFS), to test expression and efficacy *in vitro* and *in vivo* using *GBA1* L444P/L444P human induced pluripotent stem cell (iPSC)-derived dopaminergic (DA) neurons and the GD mouse model, *Gba1* D409V KI mice [44, 47, 48]. Both AAV serotypes, AAV5 and AAV9, administered via intracerebroventricular (i.c.v.) injection, achieve appropriate transduction efficiency in iPS-DA neurons and mouse brain, and rescue the GD phenotype. We further show that AAV *GBA1* delivery restores motor function in the 4L/PS-NA GD mouse model, a model characterized by low levels of GCase activity and glycosphingolipid accumulation [28]. We also tested the effects of AAV *GBA1* delivery on α-synuclein pathology in PD models *in vitro* using mouse primary neurons treated with mouse α-synuclein pre-formed fibrils (PFFs) and *in vivo,* in the CBE-chemically induced A53T M83 mouse model. In both systems, *GBA1* delivery prevents α-synuclein aggregation, a hallmark of PD.

## Materials and Methods

### Animals

#### C57BL/6 mice

C57BL/6Jcl for mouse primary neuronal cultures were purchased from CLEA Japan, Inc., Tokyo, Japan

#### Gba1 D409V KI mice

The *Gba1 D409V* mouse model (C57BL/6N-Gba1<tm1.1Mjff>/J, also known as *Gba1 D409V* KI or *Gba1 D409V/D409V*) expresses the mutant mouse D427V GCase protein, which corresponds to the D409V mutation in the mature human GCase protein [49]. Heterozygous Gba1^+/D409V^ breeders were purchased from The Jackson Laboratory, Bar Harbor, Maine, USA (Strain No. 019106). Wild-type (WT) littermates were used as control mice. A local breeding colony was established and maintained by mating heterozygous mice to produce *Gba1 D409V* KI mice at RABICS, LTD, Kanagawa, Japan.

#### A53T M83 mice

A53T α-synuclein transgenic line M83 (A53T M83) mice (B6;C3-Tg(Prnp-SNCA*A53T)83Vle/J) express mutant human A53T α-synuclein under the murine PrP promoter and harbor wild-type *Gba1* alleles [50]. Homozygous mice were purchased from The Jackson Laboratory (Strain No. 004479). A local breeding colony was established and maintained by Axcelead Drug Discovery Partners, Inc, Kanagawa, Japan.

C57BL/6 mice, *Gba1* D409V KI mice, and A53T M83 mice were bred and housed in individually ventilated cages at the Association for Assessment and Accreditation of Laboratory Animal Care (AAALAC)-accredited animal facility in Shonan Health Innovation Park, Kanagawa, Japan. Animal welfare complied with all applicable regulations, and all *in vivo* procedures described were approved by the Institutional Animal Care and Use Committee (IACUC) established in Shonan Health Innovation Park. The animal protocol numbers are as follows; Mouse primary study: AU-00030553, *Gba1* D409V KI mouse study: AU-00030684 and AU-00030616, A53T M83 mouse study: AU-00031089 and AU-00040212.

#### 4L/PS-/--NA (4L/PS-NA) mice and 4L/PS^+/+^-NA (control) mice

The transgenic GD mouse model 4L/PS-NA carry a homozygous V394L point mutation in the *Gba1* gene (4L), a complete prosaposin gene knockout (PS-) and homozygous prosaposin transgene (NA) resulting in low prosaposin levels, while control mice carry a homozygous V394L point mutation in the *Gba1* gene, but wild-type prosaposin with normal prosaposin levels [51], both of which are available at QPS Austria GmbH. Mice were bred and housed in individually ventilated cages at the AAALAC-accredited animal facility of QPS Austria GmbH [52, 53].

### iPS-derived dopaminergic (DA) neurons

For wild-type controls, the parent line 201B7 was used [54]. IPS cells with *GBA1* biallelic mutations for *L444P* were generated from control 201B7 iPS by genome editing as previously described [55]. IPS-derived DA neuronal differentiation was performed as previously described [56].

### Vector Structure and Production

Vector genome for AAV5 or AAV9-*GBA1* contains inverted terminal repeats (ITR) from AAV2, a ubiquitous promoter and intron followed by the human *GBA1* cDNA sequence, and a human growth hormone polyadenylation (hGHpA) signal sequence and a 100-bp sequence used for vector genome analysis by a quantitative polymerase chain reaction (qPCR) [57] (Patent, US 20210275614A1). The promoters used are the ubiquitous chicken β-actin hybrid (CBh) promoter with the CMV enhancer in reverse orientation (GenBank accession no. NC_006273.2) [44] for AAV-CBh-*GBA1*, and a truncated 255 bp short form of the human EF1α promoter (EFS) [47, 48] for AAV-EFS-*GBA1* (Fig 1). The vectors EFS-*GBA1* additionally contains the chimeric chicken β-actin and rabbit β-globin intron (CHIMin) [58]. Both vectors CBh-*GBA1* and EFS-*GBA1* include a 597 bp modified woodchuck hepatitis virus post-transcriptional regulatory element removing the ATGs (WPREmut6delATG) [59].

**Fig 1.**
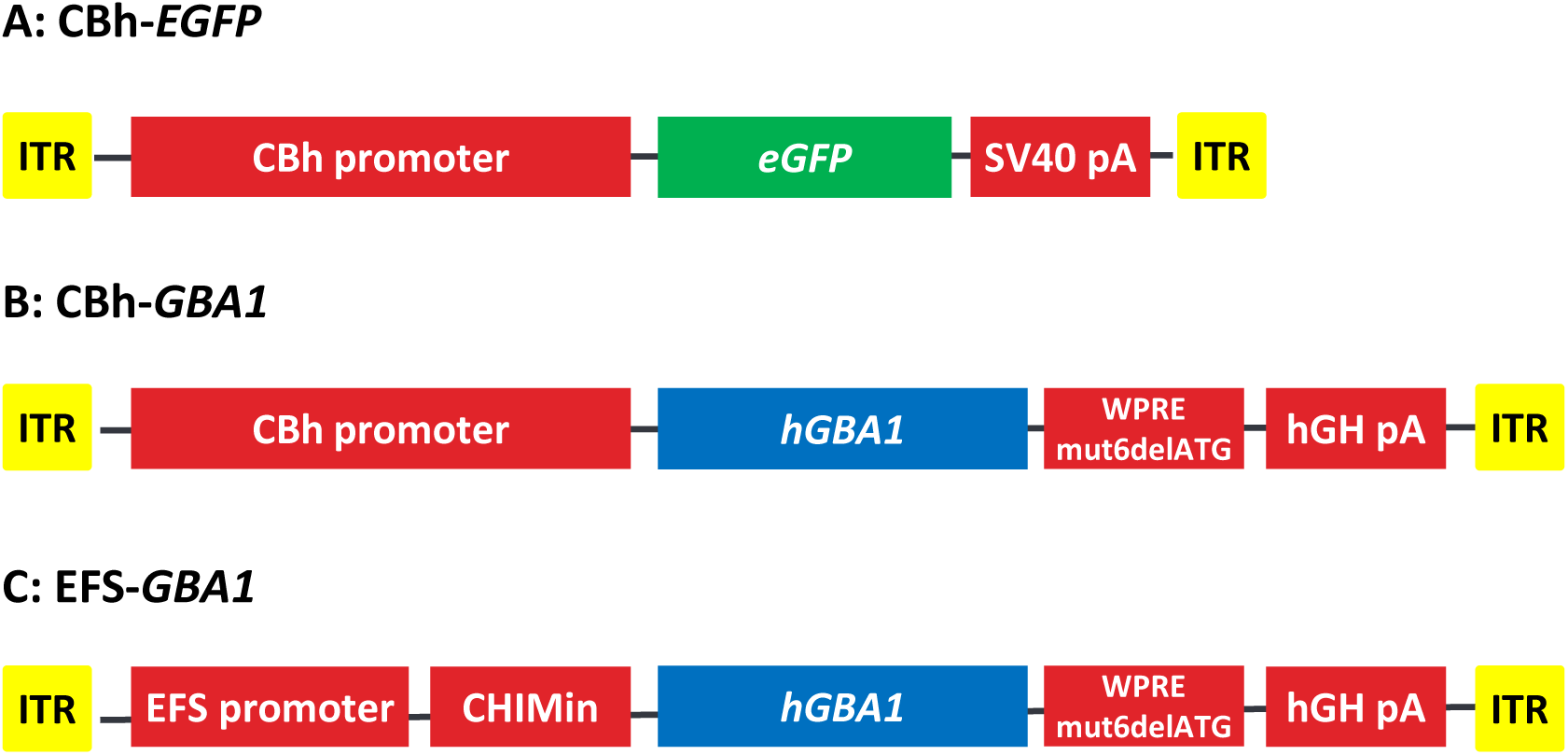
Expression cassette overview. (A) – (C) Illustration of the three expression cassettes incorporated in the AAV5 or AAV9 vectors. Abbreviations: ITR, inverted terminal repeat; WPREmut6delATG, 597 bp modified woodchuck hepatitis virus post-transcriptional regulatory element removing the ATGs; hGH poly A, human Growth hormone polyadenylation signal; CBh, chicken β-actin hybrid promoter with the CMV enhancer in reverse orientation; EFS, a truncated 255 bp short form of the human EF1α promoter; CHIMin, chimeric chicken β-actin and rabbit β-globin intron; eGFP, enhanced Green Fluorescent Protein; *hGBA1*, human *glucosylceramidase beta 1*.

The AAV5 and AAV9 vectors were produced using standard triple transfection methods. In brief, HEK293 cells were co-transfected with three plasmids: the ITR plasmid containing the expression cassette (GBA or GFP), RepCap and adenoviral helper plasmids. After 72–96 hours, the cells were chemically lysed, and the cell pellet and medium were collected. The cell lysate was cleared and treated with benzonase. The clarified lysate was loaded to a POROS CaptureSelect AAVX column connected to an AKTA™ purification system, and AAV vector-containing fractions of the elute were collected according to the manufacturer’s recommended conditions. The AAV vectors were formulated in phosphate buffered saline with 0.001% Pluronic F-68 (Thermo Fisher Scientific Inc.) and were subjected to standard characterization, including qPCR for titrating, silver staining for purity, an amebocyte lysate assay (Endosafe) for measurement of endotoxin levels (below 0.05 EU/ml). Purified vectors were aliquoted and frozen at −80°C.

### Intracerebroventricular (i.c.v.) administration of AAVs

All mice received bilateral i.c.v. injections of either vehicle phosphate buffer saline (PBS) with 0.001% Pluronic F-68) or AAV solution with stereotactic coordinates. Animals were injected bilaterally with 5 μl/hemisphere, i.e., 10 μl per brain (4L/PS-NA study) or with 10 μl/hemisphere, i.e., 20 μl per brain (*Gba1* D409V KI and A53T M83 mouse studies) at an approximate flow rate of 0.5 μl/min followed by a 5 min resting time. After anesthesia, the animals were placed into a stereotaxic apparatus, and the target regions were identified using Bregma [60] as a reference point. The following coordinates were used: for *Gba1* D409V KI and A53T M83 mouse studies, ap (anterior/posterior): −0.3 mm, ml (midline): ±1.0 mm, dv (dorsoventral): −2.0 mm. For the 4L/PS-NA mouse study the stereotactic coordinates were conducted at QPS Austria GmbH, and positions are ap (anterior/posterior): −0.5 mm, ml (midline): ±1.0 mm, dv (dorsoventral): −1.7 mm.

### Behavioral tests

The Beam Walk Test was performed three times (4, 8 and at the end of the study at 15 weeks after AAV administration) at QPS Austria GmbH. Motor coordination and balance of mice were evaluated by measuring the ability of the mice to traverse a graded series of narrow beams to reach the home cage using a 10 mm square beam for testing as previously described [61]. The testing trials were videotaped and afterwards evaluated with the Observer XT 15 (Noldus). The latency to traverse each beam and the number of slips off each beam were recorded for each trial.

### Tissue sampling

For 4L/PS-NA study, all animals were terminally anesthetized 15 weeks after AAV administration, and each sample was collected at QPS Austria GmbH. CSF was collected on the day of sacrifice by suction and capillary action until flow fully ceased in 0.2 ml polypropylene PCR tubes. For brain sampling, animals were transcardially perfused with cold 0.9% sodium chloride saline. After perfusion the brain was collected, weighed, and hemisected on a cooled surface. Then, each hemisphere of the brain was dissected in the following order: 1. Cerebellum, 2. Hippocampus, 3. Striatum, 4. cerebral cortex, and 5. midbrain including substantia nigra, snap frozen and stored at −80°C. The liver was also collected from all animals. For the GCase assay the left cerebral cortex, left hippocampus, and left striatum were used; for the GlcSph assay the right cerebral cortex, left cerebellum, CSF, and liver were used; and for mRNA/vector genome (VG) analysis the right cerebellum, right hippocampus, right striatum, right midbrain, and liver were used.

For A53T M83 mouse studies, brain samples were collected by decapitation without prior anesthesia or perfusion, followed by dissection in the following order: 1. Hippocampus, 2. Striatum, and 3. cortex, then snap frozen and stored at −80°C.

For the *Gba1* D409V KI mouse study, the brain was collected using a brain slicer. Briefly, 2-mm slices from the injection site are cut from the brain using a brain matrix. After collecting the upper 1/3 of the brain slice, the left and right sides are further separated at the midline and sampled respectively (S1 Fig).

### Quantification of vector genome (VG) and *GBA1* mRNA

DNA extraction was performed by using MagMAX™ DNA Multi-Sample Ultra 2.0 Kit (Thermo Fisher Scientific Inc.) according to manufacturer’s protocol. Briefly, DNA was extracted from tissue homogenate using the KingFisher™ Apex system (MagMAX_Ultra2_Cell_TissueCells96_v2.kfx, modified for RNase treatment). The Applied Biosystems 7500 FAST Real-Time PCR System or QuantStudio7 Flex Real-Time PCR system (Thermo Fisher Scientific Inc.) was used for quantitative PCR assay. The sequences used for the detection of tag sequence which is incorporated in each AAV expression cassette were as follows: forward primer, 5’-CCCCGTGTGAACGATTGGT-3’; reverse primer, 5’-CGTATTTCCCGTTTAGGCTTTCG-3’, and TaqMan probe, 5’-FAM-AACCCGGTGTCCTGTGAG-NFQ and MGB-3’. PCR reaction was performed for 40 cycles (95°C for 15 sec, 60°C for 1 min) after initial incubations at 50°C for 2 min and 95°C for 10 min in a volume of 20 μL containing 2× TaqPath ProAmp Master Mix, primer (150 nmol/L), probe (250 nmol/L), and water. All reactions were performed in triplicate per each sample. MagMAX™ mirVana™ Total RNA Isolation Kit (Thermo Fisher Scientific Inc.) was used for mRNA extraction. RNA extraction was performed by the KingFisher™ Apex system (MagMAX_mirVana_TissueCells_v1.kfx). Reverse transcription reaction was performed in a volume of 20 μL containing SuperScript™ IV VILO™ Master Mix (Thermo Fisher Scientific Inc.), 100 ng of mRNA template, and water to synthesize complementary DNA. PCR was performed in the same manner as for genomic DNA. The human *GBA1* sequence was detected using TaqMan Gene Expression Assays (Thermo Fisher Scientific Inc., Hs00986836_g1). The primer/probe sequences used for the detection of beta actin, an internal standard gene, was as follows: forward primer, 5’-GACTCATCGTACTCCTGCTTG -3’, reverse primer, 5’-GATTACTGCTCTGGCTCCTAG -3’, and TaqMan probe, 5’-VIC-CTGGCCTCACTGTCCACCTTCC -NFQ and MGB-3’.

### GlcSph Analysis by LC-MS/MS

For quantitation of endogenous GlcSph in iPS-DA cells, the lipid fractions were extracted by adding an ethanol and isopropanol (1:1) mixture to cells under ice-cold conditions. For quantitation of endogenous GlcSph in mouse tissues or CSF was performed using an ultra-sensitive method previously described [62], also allowing the analysis of mouse CSF. Briefly, tissue samples were homogenized in methanol to prepare 20% (w/v) homogenates for cortex and liver, and 10% (w/v) homogenates for cerebellum, hippocampus, and striatum under ice-cold conditions. Then, GlcSph from tissue homogenates was extracted by protein precipitation. For CSF samples, CSF (2 µl) was diluted with a CSF surrogate matrix (198 µl; 0.1% bovine serum albumin (BSA) and 3% heparin). GlcSph from the mixture was extracted using Strata-X polymeric solid phase extraction (Shimadzu, Kyoto, Japan). The extracts were measured by liquid chromatography tandem mass spectrometry (LC-MS/MS). The LC-MS/MS system was Shimadzu Nexera UHPLC system (Shimadzu, Kyoto, Japan) coupled to an API5000 (Applied Biosystems, CA) or QTrap6500 measured for tissue or CSF samples, respectively. Analyst software (version 1.6.3) was used for data acquisition and processing.

### GCase Activity Assay

Weighed tissues are homogenized with 10∼20 ml/g tissue homogenization buffer (250 mM Sucrose, 10 mM Tris (pH 7.5), 1 mM EDTA-2Na, 1% Triton) in Lysing Matrix D tubes by FastPrep-24 5G (MP Biomedicals) (2 cycles of 20 sec at 4 m/s with 20 sec intervals), and then supernatant was collected after centrifugation at 15,000 g for 30 min at 4°C. GCase activity was determined as described previously with some modification [63]. Briefly, GCase enzyme reaction was prepared in a clear bottom 96-well black plate by diluting a homogenate volume equivalent to 75 µg total protein with homogenization buffer to a final volume of 24.75 µl, 24.75 µl of activity assay buffer (0.25% w/v taurocholic acid in citrate/phosphate buffer) containing 2 mM 4-methylumbelliferyl-β-D-glucopyranoside substrate (4MU-Gluc; Sigma-Aldrich, # M3633) in activity assay buffer, and either 0.5 µl of DMSO or 300 mM conduritol-B-epoxide (CBE) in DMSO (final assay concentration 3 mM) were added to each sample well. The plates were incubated at 37°C for 1 h and the reaction stopped by addition of 18.75 µl of 1 M glycine (pH 10.6). 4MU fluorescence was measured using a Cytation 5 plate reader (BioTek) (excitation, 365 nm; emission, 445 nm; top read). Enzyme activity was quantified after subtracting fluorescence measured in CBE-treated (background activity) from untreated reactions. Conversion of 4-MUG to 4-MU was calculated based on a 4-MU standard curve.

### Fractionation of tissue lysates

Weighed tissues are homogenized with 10∼20 ml/g tissue homogenization buffer (20 mM Tris-HCl, pH 7.4, 50 mM NaCl, 1% Triton X-100, 0.2 mM Sodium-orthovanadate, protease inhibitor, and phosphatase inhibitor) in Lysing Matrix D tubes by FastPrep-24 5G (MP Biomedicals) (2 cycles of 20 sec at 4 m/s with 20 sec intervals. Homogenates were incubated for 30 min on ice, followed by centrifugation at 15,000 g for 60 min at 4°C. The supernatant was collected as the Triton X-100 soluble fraction. The Triton X-100 insoluble pellet was washed once with the homogenization buffer and then dissolved in the homogenization buffer containing 2% SDS using a tissue homogenizer. The resulting homogenate was centrifuged at 15,000 g for 20 min at room temperature (RT) and the supernatant was collected as the Triton X-100 insoluble fraction. Total protein concentration of Triton X-100 soluble and insoluble fraction was determined using BCA Protein Assay Kit (Thermo Fisher Scientific Inc).

### Insoluble monomeric and high molecular weight (HMW) α-synuclein detection by using ProteinSimple WES™

Human α-synuclein levels in the Triton X-100 insoluble tissue homogenate fractions were determined by using a capillary based immunoassay system (WES^TM^ ProteinSimple^®^). Samples (0.25 mg/ml protein) were applied to a 25-capillary cartridge with a 12-to-230-kDa matrix, according to the manufacturer’s protocol. After samples and antibody (anti-α-synuclein, clone 4B12, BioLegend, #807801, 1:50) had been pipetted into the pre-filled assay plate from the manufacturer, sample loading, separation, immunoprobing, washing, detection, and quantitative data analysis were performed automatically by WES™ Western System (Compass software V 4.0.0). The areas under the curve (AUC) of around 19 kDa band was used as monomeric α-synuclein quantity and AUC from 48 kDa to 230 kDa were used for high molecular weight (HMW) α-synuclein quantity, respectively.

### MSD GCase ELISA assay

GCase protein levels in the cell lysates were determined by using a customized immunosorbent assay, using MSD technology (Mesoscale discovery). Cells were lysed with 1% Triton in HEPES buffer with protease and phosphatase inhibitors, and total protein concentration was measured with the microBCA Protein Assay Kit (Thermo Fisher Scientific Inc.). 30 µl/well of capture antibody (anti-GCase antibody generated in in-house, clone 9E4), 1 µg/ml diluted in 0.2M carbonate-bicarbonate buffer) were applied to MSD MultiArray 96 well High Bind plate (MSD) and incubated overnight at 4°C. Wells were washed with wash buffer (0.05% Tween 20 in PBS) and blocked with 2% BSA (Biowest) in PBS containing 1% Casein blocker (Thermo Fisher Scientific) for 1 h at RT, with gentle shaking. After washing, 25 µl/well of cell lysates or standard (Cerezyme; ATC code A16AB02), 1:3 serial dilutions starting at 600 ng/ml to 0.82 ng/ml) were applied to the plate and incubated for 1 h at RT with gentle shaking. After washing steps, 25 µl/well of secondary antibody (anti-GCase antibody generated in in-house, clone: TK1E11-E5-G3-002) was applied conjugated with SulfoTag (1 μg/μl) and incubated for another 1 h at RT with gentle shaking. After washing and adding 150 μL MSD Read Buffer (MSD), plates were read at MSD SECTOR S600.

### Primary mouse neuron cultures

Primary mouse neurons were dissociated into single-cell suspensions from E14 mouse brains (C57BL/6Jcl, CLEA Japan, Inc.) with Neuron Dissociation Solutions (Fujifilm Wako, #291-78001) according to manufacturer’s instructions. Neurons were seeded onto 96-well poly-L-lysine-coated plates (45,000 cells/well) (Corning) and grown in Neurobasal medium (Invitrogen, #21103-049) supplemented with B-27 serum-free supplement (Invitrogen, #17504-044), GlutaMAX (Invitrogen, #35050-061), and penicillin– streptomycin in a humidified incubator at 37°C with 5% CO_2_. Half-media changes were performed once every week, or as required.

### PFF preparation and treatment in primary neurons

For treatment of neurons, α-synuclein PFFs which are provided by Prof. Hasegawa, Tokyo Metropolitan Institute of Medical Science (TMiMS) [64] (Patent, Japan 5665258JP). at concentration of 2 mg/ml was diluted in PBS at 0.2 mg/ml. They were then sonicated by Bioruputer (Diagenode) on high for 10 cycles of 30 sec on, followed by 30 sec off. The α-synuclein PFFs were then diluted in neuron media to 1.1 µg/ml and added to neuron cultures at 7 days *in vitro* (DIV). Neuron cultures were fixed and stained 12 days post-treatment.

### Immunocytochemistry

Cells were fixed with 4% (w/v) paraformaldehyde in PBS and the plate were incubated at RT for 15 min. After rinsing with PBS, cells were permeabilized with 0.1% Triton X-100 in PBS for 10–15 min followed by blocking with 5% normal goat serum (Fujifilm Wako, #143-06561) in PBS. The α-syn pSer129 antibody (1:3000, Abcam ab51253) was used as primary antibody for labeling in 1% normal goat serum in PBS and incubated overnight at 4°C. After primary antibody incubation, cells were rinsed with PBS three times, and then incubated with fluorescently labeled secondary antibodies (1:1000, Alexa594 Goat anti-Rabbit IgG, Thermo Fisher Scientific Inc., #A11037) in 1% normal goat serum in PBS containing Hoechst 33342 (1:1000, Thermo Fisher Scientific Inc., #H1399) for 1 h at RT. Automated imaging and analysis were performed with IN Cell Analyzer 6500HS (GE Healthcare). Images were acquired from 9 areas for each well. The number of Hoechst labeled cells, and numbers and intensity of pSer129 immunofluorescence staining were counted and quantified by IN Cell Developer Toolbox (GE Healthcare). The nuclear count was calculated using the Hoechst count, and pSer129 aggregation score was calculated using the pSer129 number and intensity.

### Statistics

Statistical hypothesis testing was used to compare treatment effect of control and treatment groups. When multiple treatment groups were compared to the control, Dunnett’s test was used to control its multiplicity. Adjusted p-value less than 0.05 was considered as statistically significant. Statistical analysis was performed using SAS System for Windows (Release 9, SAS Institute). The utilized statistical tests, control group to be compared, and exact sample size are mentioned in each figure legend.

Raw data (including data from behavioral tests, body weight data, and biochemical analyses) were graphed by using GraphPad Prism™ 9 (GraphPad Software Inc., USA). Graphs with group means and standard error of the mean (SEM) are provided.

## Results

### *In vitro* and *in vivo* GD phenotype was rescued by AAV5 or AAV9-*GBA1*

Human iPSCs are well-established systems for *in vitro* disease modeling. The homozygous *GBA1* L444P mutation is described to cause a severe form of GD with greatly decreased GCase levels and has been identified as PD risk gene with 5.6-fold increased risk of cognitive impairments [21, 65–67]. Our previously established iPSC-derived DA (iPS-DA) neurons carrying *GBA1* L444P biallelic mutations (L444P/L444P) [56] show reduced GCase protein levels (46.6 µg/mg protein in WT vs 6.9 µg/mg protein in L444P) and increased GlcSph accumulation (∼3 folds) (Figs 2A and 2B). We used this iPS-DA neuronal model to test recombinant AAV vectors delivering *GBA1* cDNA comparing different regulatory elements (Fig 1). The two promoters, CBh [44] and EFS promoter [47, 48] delivered by AAV9 (AAV9-CBh-*GBA1* and AAV9-EFS-*GBA1*, respectively) increase GCase expression (Fig 2A) and reduce GlcSph accumulation (Fig 2B) in a dose-dependent manner. While expression under the control of the CBh promoter plateaued at the multiplicities of infection (MOI) of 10^5^, AAV9-EFS-*GBA1* showed dose-dependent increase of expression in the dose range tested, with the highest dose (MOI of 10^6^) yielding a ∼2-fold higher GCase level than the CBh group. The increase in GCase activity correlated with reduction of GlcSph which at the highest dose tested (MOI of 10^6^) reached a ∼2-fold lower level in AAV9-EFS-*GBA1* transduced cells over AAV9-CBh-*GBA1* transduced cells (Fig 2B). To compare the transduction efficiency of AAV9 and AAV5, the CBh-*GBA1* expression cassette was used for transduction with three different MOIs. Both AAV5-CBh-*GBA1* and AAV9-CBh-*GBA1* show dose-dependent increase of GCase expression although the AAV9 vector yielded slightly higher GCase expression than the AAV5 vector at MOIs of 10^5^ and 10^6^ *in vitro*. Even at the highest dose (MOIs of 10^6^), we did not observe any obvious cell toxicity, as seen by light microscopy and further supported by the total protein concentration extracted (data not shown), while maximum GCase expression was achieved with MOI of 10^5^ (Fig 2C). In the *in vitro* iPSC system, AAV9 transduces better than AAV5, and the EFS promoter can achieve higher GCase expression.

**Fig 2.**
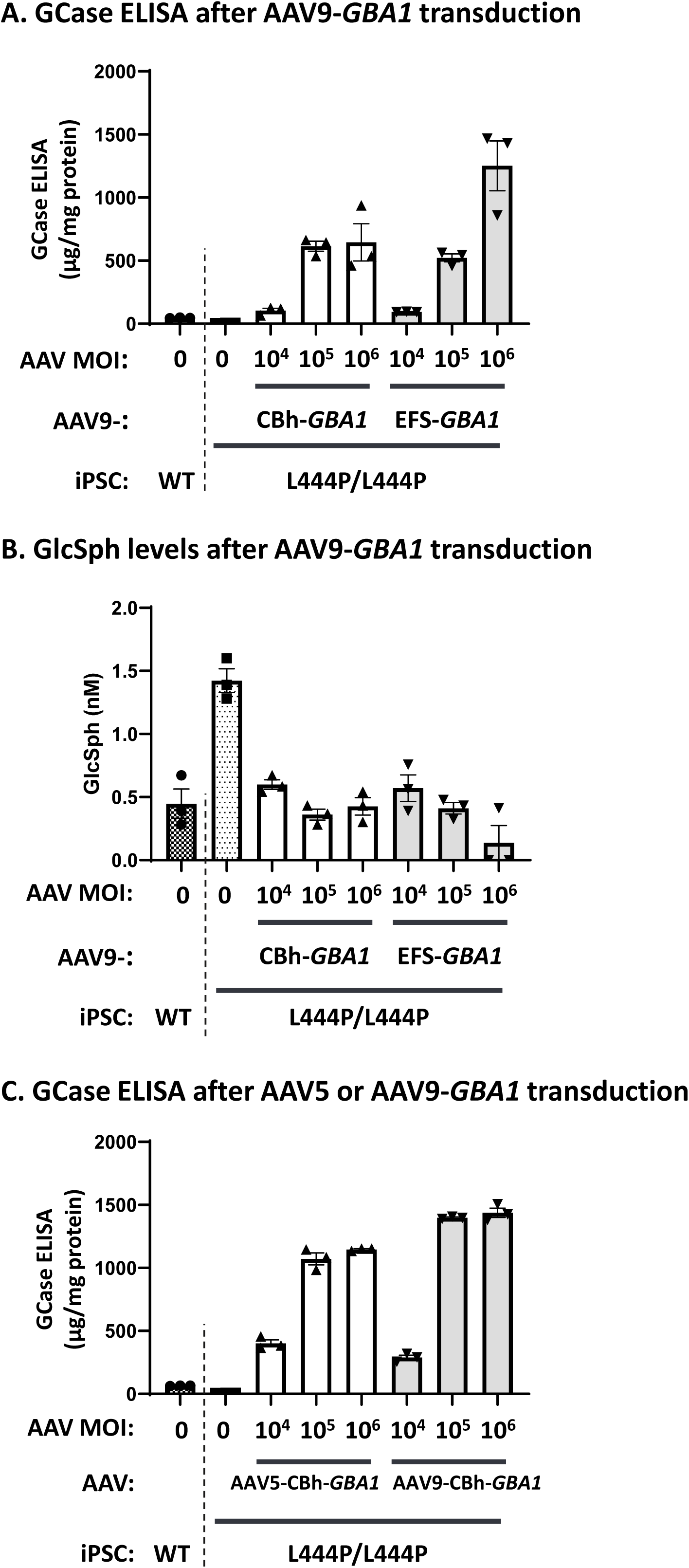
GCase expression and GlcSph reduction in iPS-derived DA neurons treated with either AAV5- or AAV9-*GBA1* vectors. Human wild-type (WT) or L444P/L444P *GBA1* mutant iPSC-derived DA neurons were transduced with AAV vectors at three MOIs (1 × 10^4^, 1 × 10^5^, and 1 × 10^6^ VG) seven days after differentiation and analyzed two weeks after AAV transduction. In (A) and (C), GCase protein levels in the cell lysate were measured using an MSD ELISA and normalized to total protein concentration assessed by BCA protein assay (mean ± SE (n = 3)). In (B), GlcSph concentration was analyzed by LC-MS/MS (mean ± SE (n = 3)). In (A) and (B), the cells were transduced with AAV9-CBh-*GBA1* or AAV9-EFS-*GBA1*, and in (C), cells were transduced with AAV5-CBh-*GBA1* or AAV9-CBh-*GBA1*.

The *Gba1* D409V KI GD mouse model, homozygous for the D409V *GBA1* mutation, was reported to have a residual GCase activity of 5% in the periphery and approximately 20% in the CNS resulting in accumulation of the glycolipids GlcCer and GlcSph [53, 68, 69]. We used this model to compare the potency of AAV9 (Study 1) and AAV5 (Study 2) carrying the same expression cassettes (CBh-*GBA1* and EFS-*GBA1*) to evaluate transduction, transgene expression, and GlcSph reduction in the mouse brain (Fig 3A). Each AAV vector was administered via bilateral i.c.v injection at a dose of 3.3 × 10^10^ vector genome (VG) per brain hemisphere (6.6 × 10^10^ VG per brain) and 4 weeks after administration GCase activity, GlcSph concentration, mRNA expression, and VG were measured in brain tissues. We detected over 40% GCase activity reduction (Fig 3B, S1 Table) and ∼60-fold increase in GlcSph accumulation (Fig 3C, S2 Table) in brain tissues from *Gba1* D409V KI mice compared to wild-type mice. Both AAV serotypes achieved significant increase of GCase activity and human *GBA1* mRNA expression with AAV9 being slightly more effective (Fig 3B, S2A Fig), and similar reduction of GlcSph accumulation in brain (Fig 3C). We detected murine *Gba1* mRNA level (2^−ΔCt^ of around 0.02) in sham treated group due to some cross-reactivity of primers with mouse *Gba1* mRNA (S2 Fig A). Even though we observed an increase of GCase activity above wild-type level (S1 Table), GlcSph reduction (3∼4 folds) (S2 Table), while highly significant, did not reach the wild-type level most likely due to AAV vector biodistribution and cell type tropism. In liver, only the AAV9 vector showed significant reduction of GlcSph accumulation (4 folds, Fig 3C, S2 Table) correlating with 50–100-fold higher VG levels measured in AAV9 compared to AAV5 treated mice (Fig 3D). When comparing transcriptional activity (*GBA1* mRNA) for the two promoters, CBh and EFS, no difference is detectable in the correlation curve between mRNA and VG, indicating that both promoters have similar strength *in vivo* (AAV5-CBh-GBA1 (Green) vs AAV5-EFS-GBA1 (Blue), S2B Fig).

**Fig 3.**
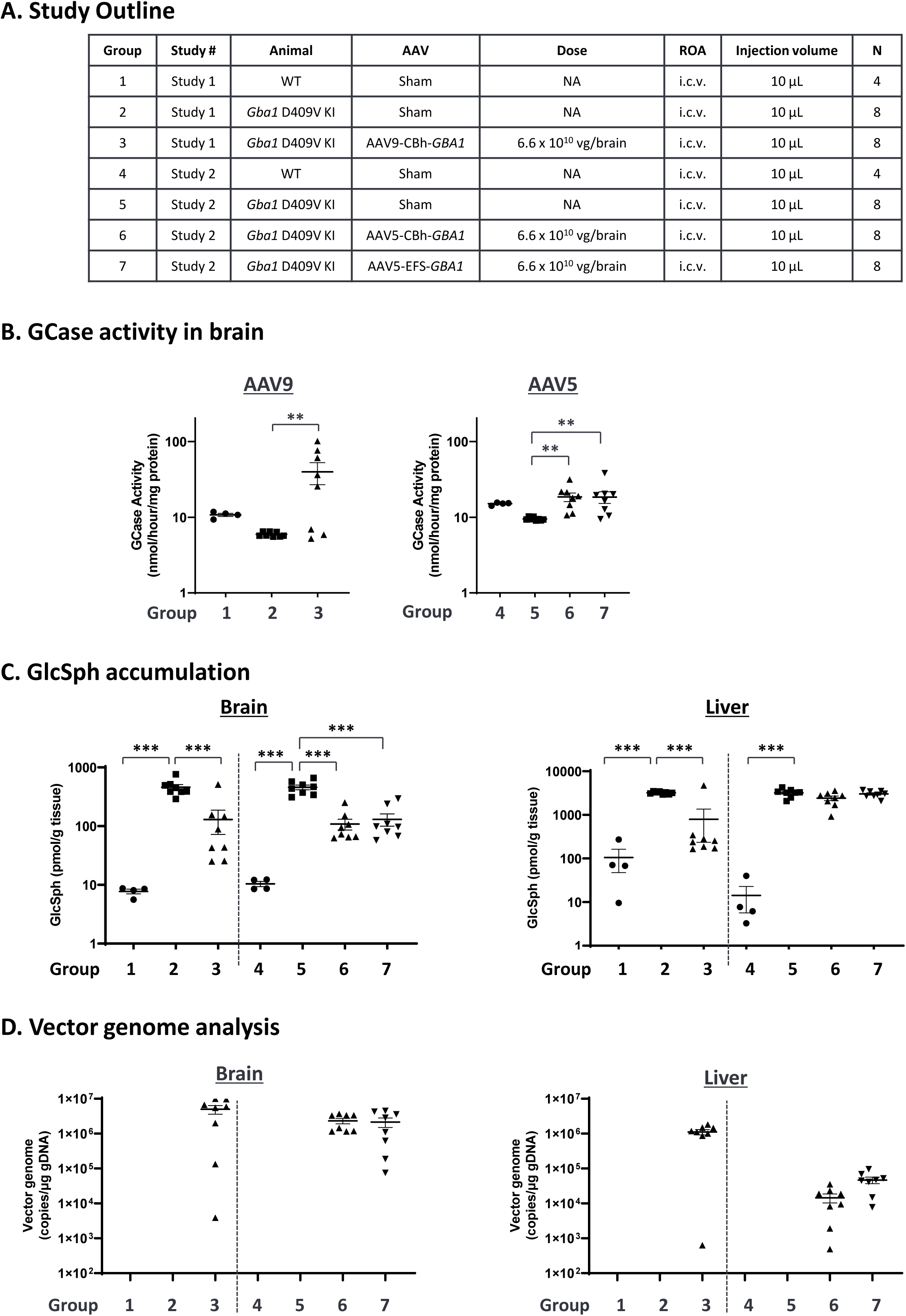
GCase activity, GlcSph, and VG analysis in *Gba1* D409V KI mouse administered with either AAV5 or AAV9-*GBA1*. The study outline is described in the table (A). AAV5-*GBA1* or AAV9-*GBA1* were delivered by bilateral i.c.v. injection of buffer (sham) or AAV (3.3 × 10^10^ VG/ hemisphere, total 6.6 × 10^10^ VG/brain) in 16-week-old male *Gba1* D409V KI mice or WT littermates. Mice were sacrificed 4 weeks after injection. Brains were isolated and sliced as 2-mm-thick brain slices around the injection site, and tissues were analyzed for GCase enzymatic activity, GlcSph level, and VG. (B) GCase activity was measured in each independent study (Study 1 and Study 2). See S1 Table for mean values. (C) GlcSph concentration was analyzed by LC-MS/MS. See S2 Table for the mean values per group. (D) VG in brain and liver was measured by qPCR specific for DNA detection. VG in the liver and brain of groups 1, 2, 4, and 5 were below lower limit of quantitation (LLOQ) as expected as these were sham controls. For B, C and D, each graph represents mean ± S.E.M. (n = 4 or 8) with the Y-axis as logarithmic scale. For B and C, statistical analyses were performed by Dunnett analysis. *: <0.05; **: <0.01; ***: <0.001, compared to Group 2 and Group 5 for batch 1 study and batch 2 study, respectively. For D, statistical analysis is not performed since the control value is zero.

### Recovery of motor performance in 4L/PS-NA mouse model by AAV5-EFS-*GBA1*

The GD mouse model, 4L/PS-NA is characterized by strong motor deficits, low levels of prosaposin and saposin expression and functionally impaired GCase, due to the homozygous V394L point mutation in the mouse *GBA1* gene [51–53]. Two AAV5-*GBA1* vectors, AAV5-CBh-*GBA1* and AAV5-EFS-*GBA1*, were administered at a dose of 2.7 × 10^10^ VG per brain hemisphere (5.4 × 10^10^ VG per brain) into 4-week-old mice by bilateral i.c.v. (Fig 4A). Mice from all groups showed normal body weight gain during the study, indicating that the treatment was well tolerated (S3A Fig). As the 4L/PS-NA model represents a severe disease model, we lost three animals during the study for reasons unrelated to the procedures or AAV administration, one in Group 2 (Sham) and two in Group 3 (AAV5-CBh-*GBA1* group) (Fig. 4A). Several brain regions were isolated and assigned to either GCase activity, GlcSph detection, or VG and mRNA profiling. GCase activity was measured in cortex, hippocampus, and striatum, and GlcSph substrate levels were analyzed in cortex, cerebellum, CSF, and liver. Following AAV5-*GBA1* treatment, GCase activity increased in cortex (3.1 and 2.0 folds in CBh and EFS promoter, respectively) and hippocampus (54 and 64 folds in CBh and EFS promoter, respectively), and striatum (1.9 and 1.3 folds in CBh and EFS promoter, respectively) (Fig 4B, S3 Table). The highest GCase activity was measured in hippocampus probably due to the higher biodistribution of AAV5 into this brain region after i.c.v. which is supported by the VG data (S3B Fig). Remarkably high GlcSph accumulation was detected in brain, liver, and CSF of 4L/PS-NA mice compared to control mice (Fig 4C). 4L/PS-NA mice have a significant increase of GlcSph accumulation while having similar GCase activity level to that of control mice, which can be explained by the fact that saposin C, a cleavage product of prosaposine and cofactor of GCase, is required for GCase activity *in vivo* but is dispensable for the *in vitro* assay used to measure GCase activity in tissue homogenates. Administration of AAV5-*GBA1* via i.c.v. resulted in significant substrate reduction in cortex (2.2-fold reduction for both CBh-*GBA1* and EFS-*GBA1*), cerebellum (1.7- and 1.3-fold reduction for CBh-*GBA1* and EFS-*GBA1*, respectively), and CSF (2.1- and 1.6-fold reduction for CBh-*GBA1* and EFS-*GBA1*, respectively), while no substrate reduction was detected in liver (Fig 4C, S4 Table), consistent with the data shown above demonstrating limited VG distribution for AAV5 to the liver after i.c.v. administration (Fig 4C and S3C Fig). In this study, we were able to detect GlcSph in a small volume of CSF using a highly sensitive quantitation method established for human CSF samples [62]. In summary AAV5-delivered *GBA1*, independently of the expression cassette used (AAV5-CBh-*GBA1* or AAV5-EFS-*GBA1*), increases GCase activity and reduces GlcSph accumulation in the brain of 4L/PS-NA mice.

**Fig 4.**
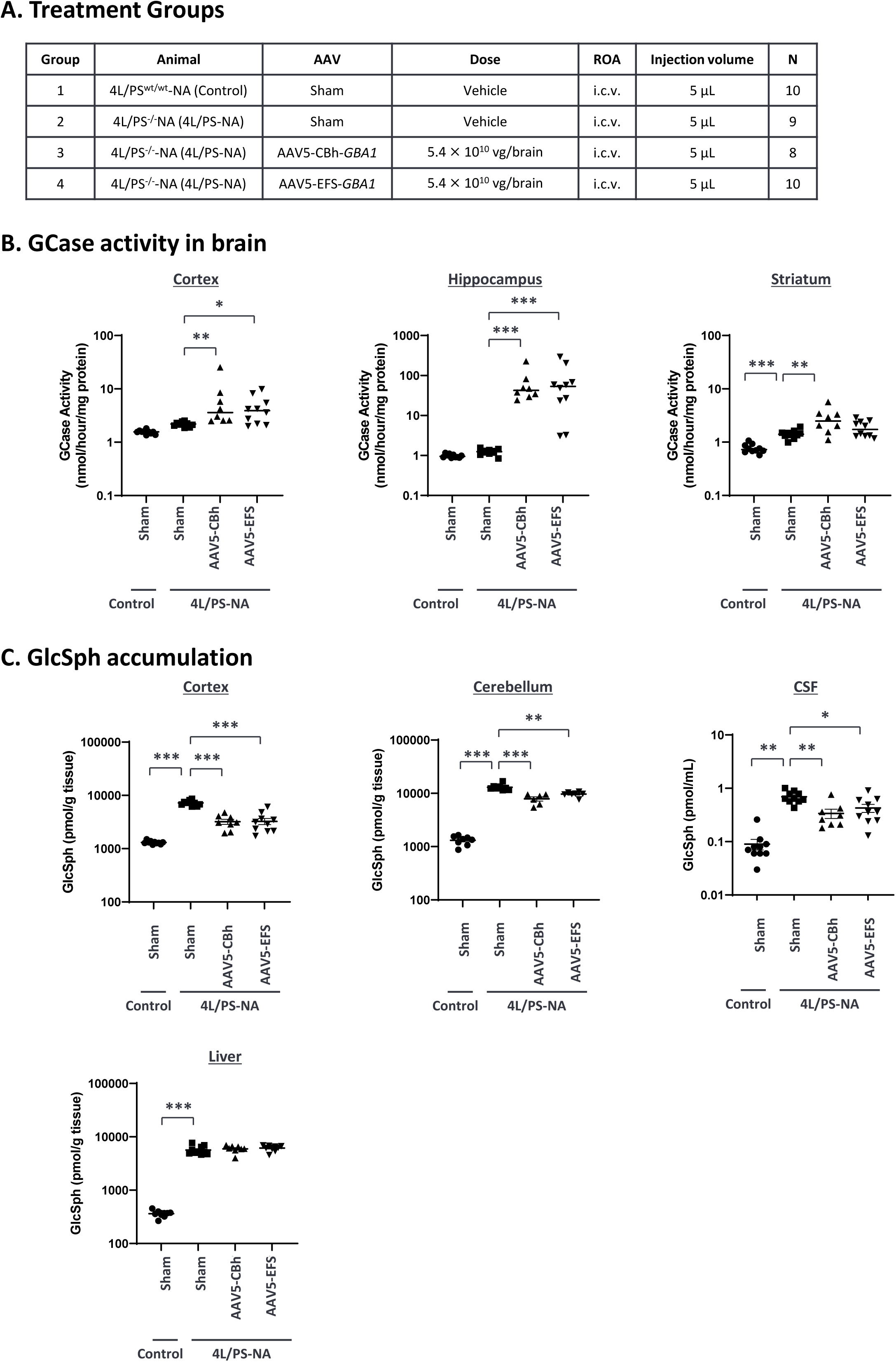
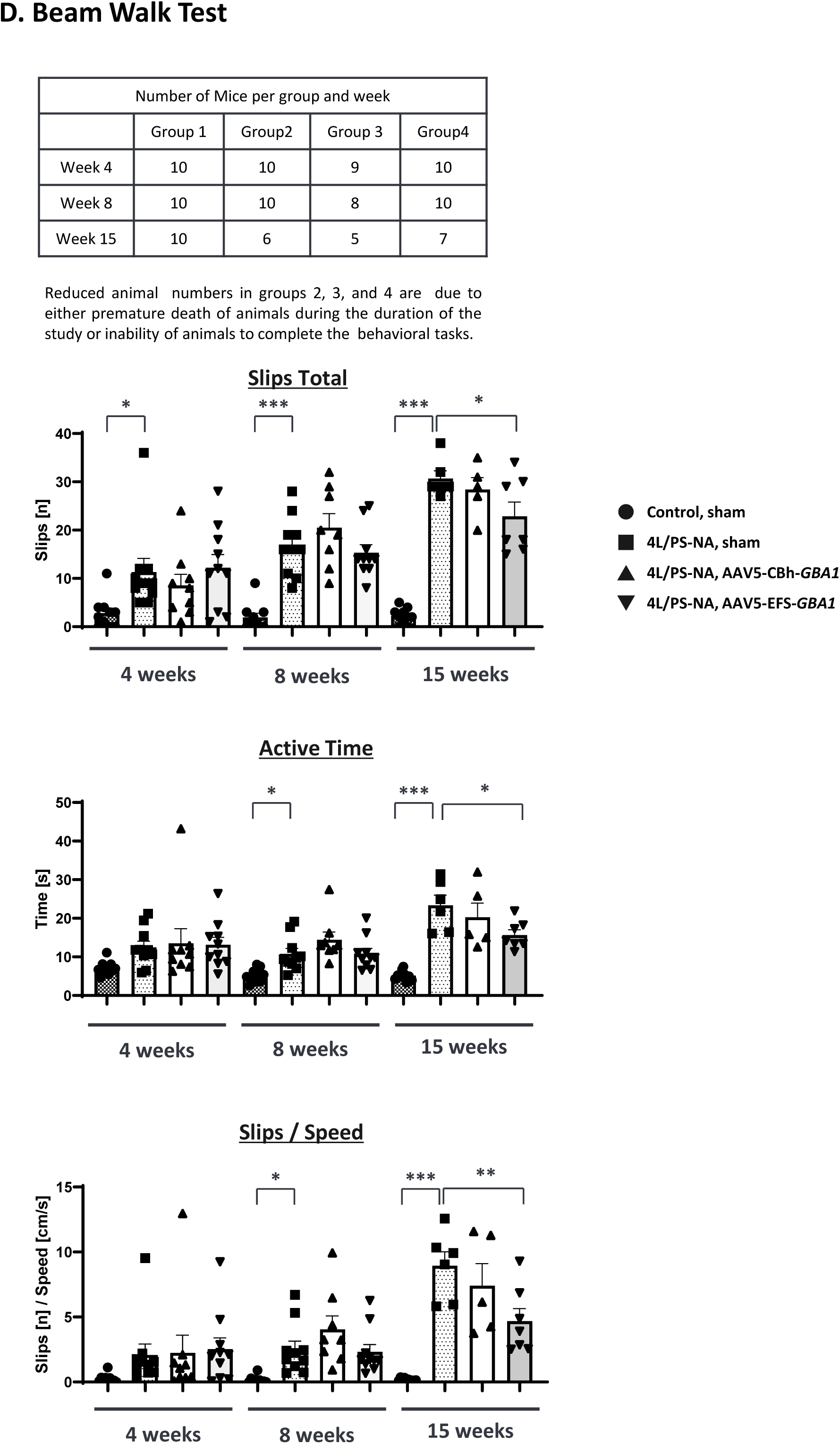
Biochemical and behavior analysis in 4L/PS-NA mouse model treated with AAV5-*GBA1*. Table. (A) illustrates the study overview. 4-week-old male 4L/PS^wt/wt^-NA (control) and 4L/PS^−/−^-NA (4L/PS-NA) mice were injected with buffer (sham), AAV5-CBh-*GBA1* or AAV5-EFS-*GBA1* (i.c.v., bilaterally, 2.7 ×10^10^ VG/hemisphere, total 5.4 × 10^10^ VG/brain). After 15 weeks, brain and liver tissues were isolated. In Group 2 one mouse died prematurely and in Group 3 two mice died prematurely, reducing the group size to 9 and 8, respectively. (B) GCase activity was measured in three different brain regions. The Y-axis is logarithmic scale. The stars indicate the statistical difference compared to the control Group 2. See S3 table for the mean values for GCase activity. (C) GlcSph concentration was measured by using MS/MS analysis. The Y-axis is logarithmic scale. The stars indicate the statistical difference compared to the control Group 2. See S4 table for the mean values for GlcSph concentration. (D) The Beam Walk Test was performed three times (at 4, 8 and 15 weeks after AAV administration). Graphs represent total slips [n], active time [s], total slips [n] per speed [cm/s] for each group on the beams tested at all time points. Data are shown as means ± SEM per group. Statistical analyses were performed by Dunnett analysis. *: <0.05; **: <0.01; ***: <0.001 compared to Group 2 (4L/PS-NA, sham group) in each timepoint.

To assess the effect of AAV5-delivered *GBA1* on motor performance, the beam walk test was conducted three times (4, 8, and 15 weeks after AAV administration). 4L/PS-NA animals exhibited significantly reduced performance in all readouts and at all time points tested. 4L/PS-NA mice treated with AAV5-EFS-*GBA1* (group 4) but not AAV5-CBh-*GBA1* (group 3) showed significantly improved motor performance 15 weeks after AAV administration, indicated by decreased slip total numbers, active time, and slips/speed, when compared to 4L/PS-NA sham mice (group 2) (Fig 4D). These results show that AAV5-EFS-*GBA1* delivery to the brain improves motor deficits in the 4L/PS-NA GD mouse model.

### Rescue of α-synuclein phenotype in an *in vitro* mouse primary neuron model by AAV9-*GBA1*

Given the association between reduced GCase activity and the accumulation of α-synuclein in PD, our study aimed to determine the impact of GCase on modulating α-synuclein pathologies. We have used a mouse primary neuronal *in vitro* model for α-synuclein aggregation designed to mimic PD pathology [70–72]. First, the dependence of phospho-α-synuclein at Ser 129 (pSer129-α-Syn) on the dose of α-synuclein PFFs and duration of treatment was established (data not shown), and optimal conditions were selected (1.1 ng/ml PFFs and 12-day culture; Fig 5A). Using this model, we assessed whether increasing expression of GCase alters the pSer129-α-Syn pathology. Two AAV9-*GBA1* vectors (CBh-*GBA1* and EFS-*GBA1*), as well as the negative control AAV9-CBh-*EGFP* were tested at three different MOIs, 1 × 10^3^, 1 × 10^4^, and 1 × 10^5^. After 12 days of treatment, we observed EGFP expression in approximately 20%, 50% and over 90% of cells at the MOI of 1 × 10^3^, 1 × 10^4^, and 1 × 10^5^, respectively (S4A Fig), though the nuclear number was reduced at the highest MOI (Fig 5B), which may indicate a dose dependent AAV9 toxicity. At the lowest MOI (1 ×10^3^), no significant difference was observed between groups. At a dose of 1 × 10^4^ and 1 × 10^5^, the AAV9-*GBA1* groups using either vector, AAV9-CBh-*GBA1* or AAV9-EFS-*GBA1*, resulted in reduced pSer129-α-Syn aggregation for both criteria, number of pSer129-α-Syn aggregates and overall signal intensity, with AAV9-EFS-*GBA1* performing slightly better than AAV9-CBh-*GBA1*. At the highest dose of AAV9-EFS-*GBA1* a 6∼8-fold reduction of pSer129-α-Syn granules was detected. The AAV9-*EGFP* did not show any effects on pSer129-α-Syn aggregation (images in S4A and S4B Figs and graphs in Figs 5C and 5D). These results demonstrate that *GBA1* over-expression effectively resolves pSer129-α-Syn aggregates, confirming the involvement of GCase expression in the clearance of α-synuclein pathology.

**Fig 5.**
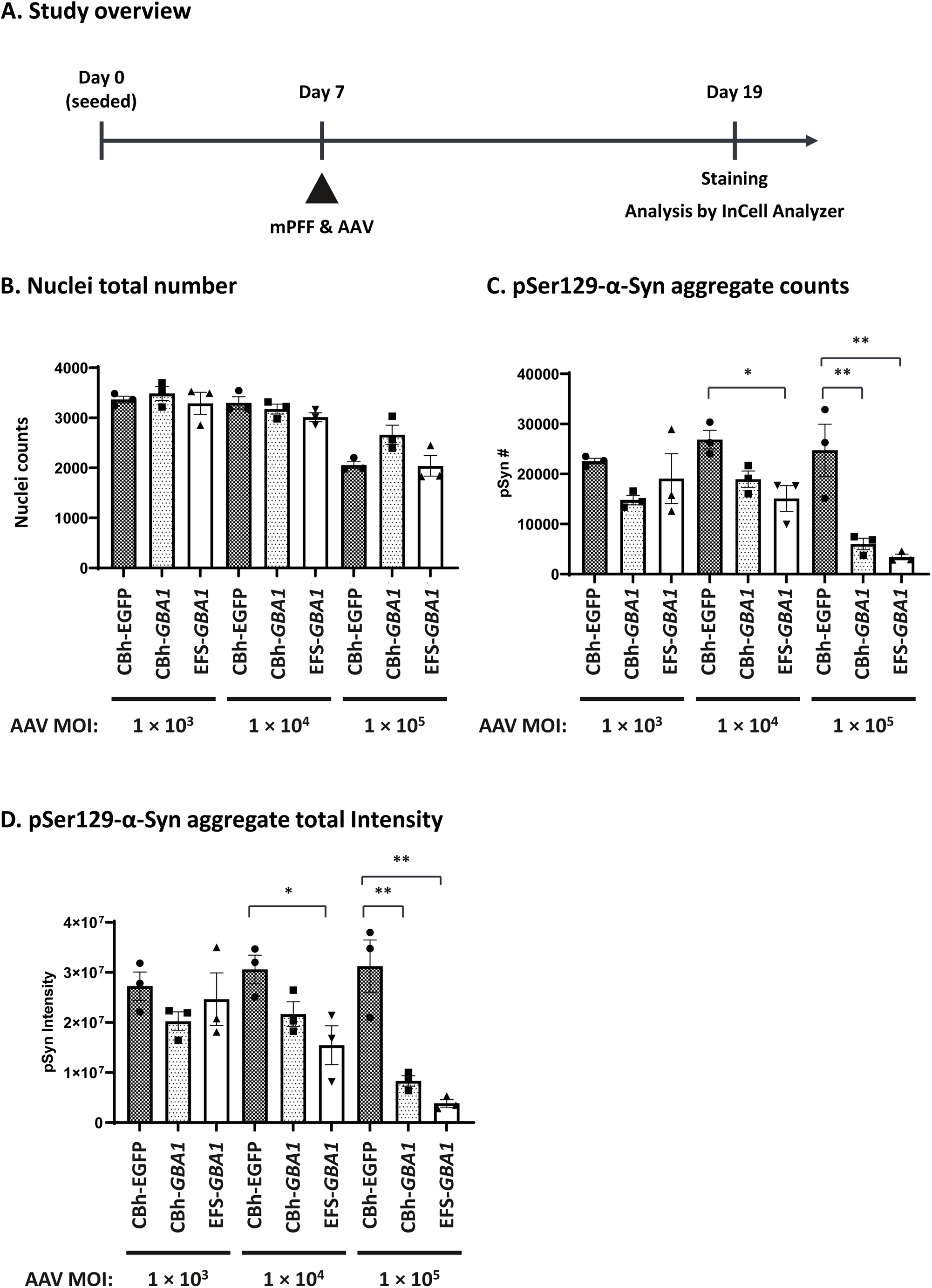
Phospho-α-synuclein reduction in mouse primary neuron cells treated with AAV9-*GBA1*. (A) illustrates the study layout. Mouse primary cortical neurons were seeded on poly-D-Lysin coated 96-well plate (45,000 cells/well) on Day 0. On Day 7, mouse pre-formed fibril (mPFF; 1.1 ng/ml) and AAV9-*EGFP* or AAV9*-GBA1* (MOI at 1 ×10^3^, 1 × 10^4^, or 1 × 10^3^ VG) were added to the cells as noted on the x-axis. After 12-day culture, cells were stained with anti-phospho-α-synuclein antibody and analyzed by InCell Analyzer. Quantification of the nuclei counts is shown in (B), the pSer129 α-Syn aggregate counts in (C), and the intensity of pSer129 α-Syn aggregates in (D). Data are represented as mean ± S.E.M. (n = 3). Statistical analyses were performed by Dunnett analysis. *: <0.05; **: <0.01; ***: <0.001 compared to CBh-*EGFP* control in each dose (MOI).

### Rescue of α-synuclein phenotype in an *in vivo* mouse model by AAV5-*GBA1*

Though the relationship between GCase reduction and α-synuclein pathology has been previously described, the phenotypes of the published models tend to develop slowly and lack robustness [31, 32, 73, 74]. To establish a model which shows faster and more robust *GBA1*-associated α-synuclein pathology, we utilized the combination of the chemical GCase inhibitor CBE in the genetic engineered A53T M83 mouse model [50, 74].

To select the appropriate CBE dose, we performed a dose-finding study using 1, 5, and 25 mg/kg of CBE with intraperitoneal (i.p.) injection in the A53T M83 mice, and measured body weight, GCase activity, and GlcSph levels (S5 Fig). Clear dose-dependent GCase activity reduction (S5C Fig, S7 Table) and accumulation of GlcSph (S5D Fig, S8 Table) were observed without any changes in body weight or other signs of gross toxicity (S5B Fig). We selected the 25 mg/kg CBE regimen for subsequent experiments as this dose shows a clear change in both GCase activity and GlcSph, without reaching saturation and is anticipated to elicit change in other GCase-dependent phenotypes.

Next, to determine if GCase inhibition causes α-synuclein pathology and if GCase supplementation can restore the phenotypes, CBE was administered via i.p. injection at 25 mg/kg once daily for 10 days in the A53T M83 mice two weeks after AAV5-EFS-*GBA1* was injected bilaterally via i.c.v. with 1.7 × 10^10^ VG per brain hemisphere (3.4 × 10^10^ VG/brain) (Fig. 6A). Brain tissues (cortex, hippocampus, and striatum) were collected and analyzed for GCase activity, GlcSph levels, and α-synuclein pathology, which is the high-molecular-weight α-synuclein (HMW α-Syn) in Triton X-100 insoluble fraction. Mice from all groups showed normal body weight gain during the study indicating that the treatment was well tolerated (S6A Fig). In the striatum, GCase activity was significantly reduced after CBE treatment, and could be rescued by AAV5-EFS-*GBA1* administration, thereby restoring GCase activity to wild-type levels (2.9-fold increase) (Fig 6B, S5 Table). GlcSph measurements in the striatum showed significantly increased GlcSph levels after CBE treatment (∼100 folds compared to control), which was significantly reduced after AAV5-EFS-*GBA1* administration (1.7-fold reduction) (Fig 6C, S6 Table).

**Fig 6.**
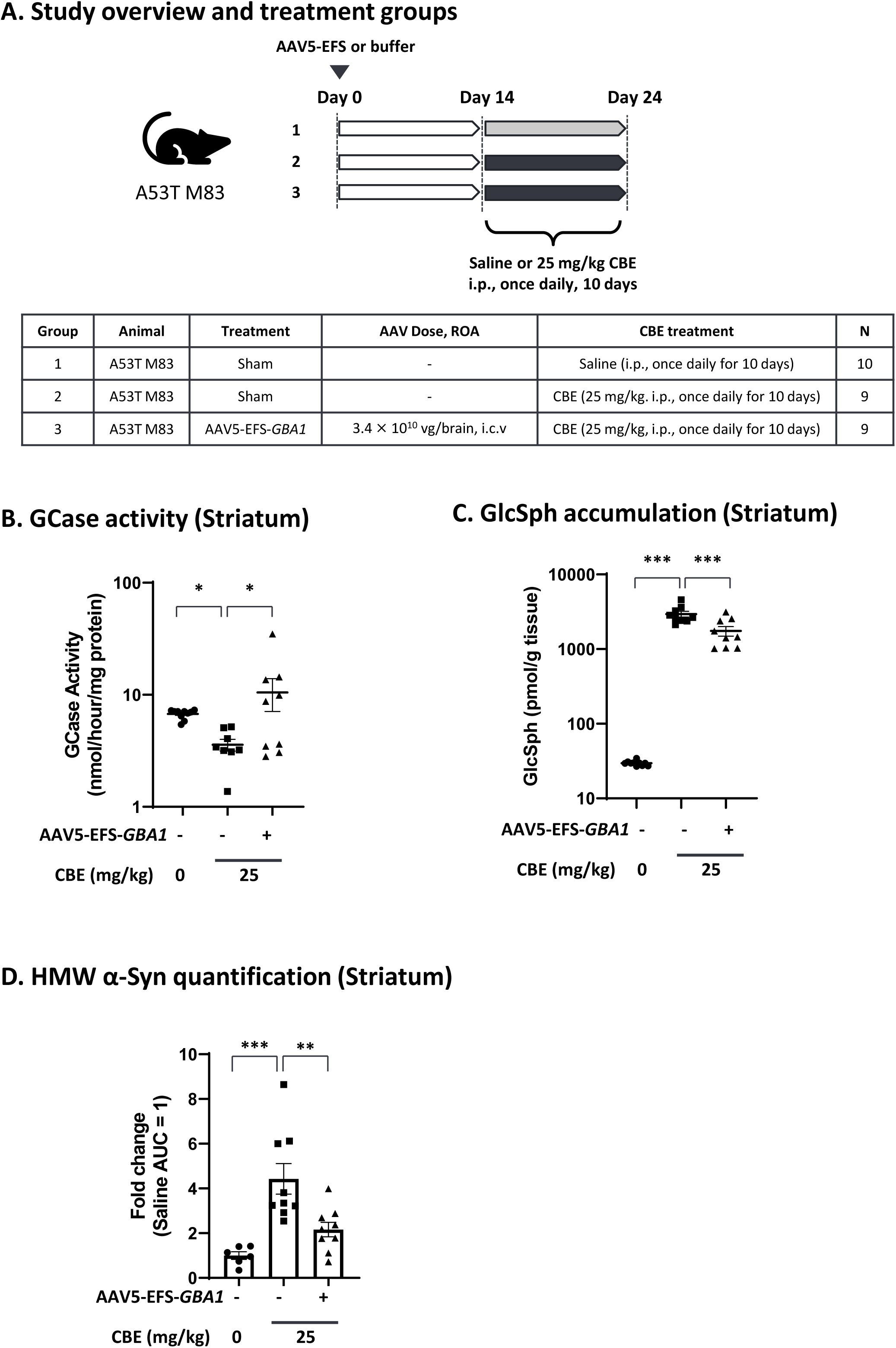
Efficacy of AAV5-*GBA1* on GCase activity, GlcSph accumulation and HMW α-synuclein in A53T M83 mouse model treated with CBE. In (A), the study outline is illustrated. 12-week-old male A53T M83 mice were administered with AAV5-EFS-*GBA1* (i.c.v., bilaterally, 1.7 ×10^10^ VG/hemisphere, total 3.4 × 10^10^ VG/brain) on day 0. For Group 2 and 3 CBE treatment (25 mg/kg, i.p., once daily) started two weeks (day 14) after AAV administration and continued for 10 days (up to day 24). At the end of the study (day 24) striatum was analyzed for GCase activity (B), stars indicate statistical significance over the control Group 2. (C) shows GlcSph levels, stars indicate statistical significance over the control Group 2. In (D) the AUC was normalized with the average of Group 1 quantified by WES™. See S5 and S6 tables for mean values for GCase activity and GlcSph levels. Statistical analyses were performed by Dunnett analysis. *: <0.05; **: <0.01; ***: <0.001 compared to group 2 (sham, CBE).

We analyzed aggregated α-synuclein by quantifying both monomeric and HMW α-Syn in striatum, hippocampus, and cortex. A significant HMW α-Syn increase was observed only in the striatum (Fig 6D, and S6B Fig), but not in the cortex and hippocampus, possibly due to the high baseline of HMW a-Syn observed in control animals not treated with CBE (data not shown). In the striatum, HMW α-Syn was about 50% reduced after AAV5-EFS-*GBA1* treatment, however, still about 2-fold above untreated control levels (Fig 6D and S6B Fig), while monomeric α-Syn did not change (S6C Fig). These data suggest that GCase inhibition with CBE increases HMW α-Syn accumulation, one of the key PD pathologies, and that such can be rescued by *GBA1* overexpression.

## Discussion

In this study, we have evaluated two AAV serotypes, AAV5 and AAV9, and two ubiquitous promoters, CBh and EFS, to better understand their effectiveness both *in vitro* and *in vivo*, in iPS-DA neurons and mouse models. We have demonstrated that GCase overexpression restores not only GD phenotypes including GlcSph accumulation and motor dysfunction, but also the critical PD phenotype, α-synuclein pathology, both *in vitro,* in iPS-DA neurons or mouse primary neurons and *in vivo,* in mouse disease models for both GD and PD.

When we compared the two promoters, CBh and EFS, in neuronal cells from iPS-DA neuron cultures, we found that AAV9-CBh-*GBA1* reaches saturated expression at mid and the high dose (MOI of 10^5^ and 10^6^) whereas AAV9-EFS*-GBA1* shows a dose-dependent increase of expression to higher GCase levels, and slightly higher reduction of GlcSph at the highest dose (Figs 2A and 2B). In agreement with this finding, AAV9-EFS*-GBA1* performs somewhat better than AAV9-CBh-*GBA1* for the rescue of the pSer129-α-Syn phenotype in mouse primary neurons (Figs 5C and 5D). This may indicate that EFS is more active in neuronal cells compared to CBh. In two mouse models, *Gba1* D409V KI mice (Figs 3B and 3C) and 4L/PS-NA mice (Figs 4B and 4C), both promoters perform comparable for GCase activity and GlcSph reduction. The correlations between VG and human *GBA1* mRNA expression suggest that the strength of the two promoters is similar (S2B Fig, green vs blue dots; S3D Fig, open vs closed circles). Interestingly, however, only AAV5-EFS*-GBA1* rescues motor behavior in the 4L/PS-NA model (Fig 4D). This difference may correlate with the observation that EFS performs better in *in vitro* neuronal cell cultures and may also have higher activity in neuronal cells *in vivo.* The effect may be due to EFS having a slightly higher activity surpassing the threshold required for rescue of motor activity, or that EFS activity is higher in specific cell types vs CBh, as suggested by the *in vitro* data (Fig 2A, Figs 5C and 5D). These data suggest that it is important to include multiple parameters to assess promoter functionality in order to choose the appropriate expression construct, especially when developing a therapeutic treatment. Here we see clear differences in the *in vitro* iPSC system and in the motor behavior in the 4L/PS-NA mouse model.

We next compared AAV5 and AAV9 transduction efficiency in the iPS-DA neuron model and mouse brain. In iPS-DA neurons we used GCase protein expression as readout and demonstrated that AAV9 performs better at the MOI of 10^5^ and 10^6^ (1.3-fold increase over AAV5) (Fig 2C). In general, we noticed that AAV9 works more consistently *in vitro,* and therefore is preferred for *in vitro* evaluation. We compared AAV5-CBh-*GBA1* and AAV9-CBh-*GBA1* biodistribution in the *Gba1* D409V KI mouse model after i.c.v. administration. For GCase activity and *hGBA1* mRNA we see more variability for AAV9 compared to the AAV5 group, and overall higher expression for the AAV9 group (Fig 3B, S2A Fig). Both vectors achieved the goal to restore at least wild-type levels of GCase. Assessing the downstream effect, the reduction of GlcSph in brain (Fig 3C, left), comparable levels between AAV5 and AAV9 are observed. A major difference between AAV5 and AAV9 is seen when GlcSph levels are analyzed in the liver, where only the AAV9 group shows reduction of GlcSph levels (Fig 3C, right). This finding correlates with the AAV VG data (Fig 3D), where only AAV9 shows significant transduction of the liver, which further correlates with the h*GBA1* mRNA expression (S2A Fig).

These data show a clear difference between AAV9 and AAV5, regarding liver tropism after i.c.v. administration, with a significant leakage to the periphery in the case of AAV9. For AAV5 the distribution to the liver is minimal, which would be beneficial when delivery to brain only is favored, and it would avoid the potential hepatotoxicity, a known liability of AAVs [75]. Still, AAV9 may be preferred for indications like GD, where both CNS and peripheral symptoms need to be treated. Further validation in large animals such as non-human primate (NHP) is necessary to confirm the tropism since differences have been reported between rodents and NHPs, and old-world monkeys are currently the gold standard for translation to human [76, 77]. In conclusion, we selected AAV9 for our *in vitro* model based on the better transduction efficiency, and AAV5 for our *in vivo* model due to the low tropism to liver and reduced risk of hepatoxicity.

To examine if AAV5-*GBA1* can rescue GD phenotypes, we tested two promoters in the neuropathic GD mouse model 4L/PS-NA [52, 78]. At the doses used, we did not observe any gross abnormalities due to the AAV administration as demonstrated by normal body weight gain (S3A Fig). Our results show that both AAV5-*GBA1* expression cassettes (CBh-*GBA1* and EFS-*GBA1*) significantly reduce the GlcSph accumulation in cortex, cerebellum, and CSF, with no difference between the promoters (Fig 4C). GCase expression levels were similar for both promoters, with hippocampus showing the highest expression levels, as well as the highest VG (Fig 4B and S3B Fig). This finding suggests that i.c.v. administration in mice achieves efficient delivery to the hippocampus, which may be specific for mice and will need to be further evaluated in large animals to understand the translational impact. As already discussed above there is exceptionally low transduction of AAV5 in the liver, and consequently no *hGBA1* mRNA expression and no effect on GlcSph levels. Next, we performed the Beam Walk Test to assess if AAV-*GBA1* transduction can restore the dysfunction of motor coordination and balance of animals. 4L/PS-NA mice showed phenotypes already at 4 weeks after surgery compared to control animals which worsened over time. We performed the test at three timepoints and found that AAV5-EFS-*GBA1* could rescue the phenotypes only at the last timepoint (15 weeks after AAV administration). One possible explanation is that since this assay has a big variation, it only shows significant difference with the big window of parameters between 4L/PS-NA and control mice, or it may take time to show the efficacy after GCase expression.

The accumulation and aggregation of α-synuclein plays a major role in PD, with α-synuclein fibrils being toxic causing neuronal death [79]. The link between GCase function and α-synuclein pathology has been proposed from post-mortem tissue analysis showing an inverse correlation of low GCase and high α-synuclein pathology [14, 80, 81]. Several *in vitro* studies also show that this inverse correlation and that GCase activity is inhibited by α-synuclein fibrils [82–84]. To understand if there is a direct interaction between GCase activity and α-synuclein pathology, GCase was inhibited by shRNA knockdown or CBE treatment and α-synuclein aggregation in response to α-synuclein PFFs treatment in primary neuron cells was evaluated, showing that GCase inhibition increased phospho-α-synuclein aggregation and/or release of α-synuclein fibrils [70, 71, 82]. However, whether GCase supplementation can sufficiently revert the α-synuclein phenotype remains unclear. Thus, we tested the effect of AAV-*GBA1* on phospho-α-synuclein aggregation induced by PFF addition in mouse primary neuron cells and demonstrated that these α-synuclein aggregations are dissolved by *GBA1* overexpression (Figs 5C and 5D). This is the first evidence demonstrating the direct effect of GCase on α-synuclein aggregates *in vitro,* and this *in vitro* system is useful to elucidate how GCase activity plays a role in α-synuclein pathologies and can be used as a screening tool for GCase modulators in neural cells.

To examine the relationship between GCase on α-synuclein pathology *in vivo*, we used the A53T M83 mouse model to evaluate the effect of GCase on α-synuclein pathology in the mouse brain by measuring the aggregated form of α-synuclein, detectable as HMW (oligomer) forms in Triton X-100-insoluble fraction. After CBE treatment the amount of HMW α-Syn accumulation rises by about 4-folds in the striatum, confirming the inverse correlation of GCase activity and α-synuclein pathology. In the group treated with AAV5-*GBA1* before CBE treatment HMW α-Syn accumulation was greatly reduced (Fig 6D). Interestingly, in cortex and hippocampus we could not detect the increase of HMW α-Synuclein (data not shown). While we do not have a clear explanation, this fits with the observation that substantia nigra is more sensitive to α-synuclein aggregation and it may need longer CBE treatment to mimic the phenotypes in cortex and hippocampus [74]. Previously, it has been reported that a single injection of CBE increases α-synuclein levels limited to the substantia nigra, and does not affect cortex or hippocampus, suggesting different sensitivities of CBE in different brain regions [85]. Another possibility is different susceptibility to α-synucleinopathy in different brain regions or in cell types as described in the past [86, 87]. These data indicate that increasing GCase activity in mouse brain can restore cellular homeostasis of degrading α-synuclein aggregation and therefore the potential to slow disease progression caused by α-synuclein aggregation.

We have demonstrated that GCase overexpression can decrease the levels of the oligomeric and aggregated forms of α-synuclein (HMW insoluble α-Syn and/or pSer129-α-Syn). Our *in vivo* model demonstrates that GCase reduction by CBE treatment is sufficient to induce aggregation of α-synuclein in the striatum, one of the most affected regions in PD. In the neuronal *in vitro* cell system GCase supplementation can revert α-synuclein aggregation induced by PFFs, suggesting that GCase expression alone can be effective to treat α-synucleinopathies independent of a *GBA1* mutation status. In idiopathic PD patients reduced GCase activity has also been reported independent from the *GBA1* mutation status and it is thought that lysosomal dysfunction and age-related diminished lysosomal capacity contribute to the accumulation of α-synuclein [81, 82, 88]. It has also been reported that GCase deficient cells increases the release of α-synuclein both *in vitro* and *in vivo* [82, 89], suggesting that GCase is a key factor for the vicious cycle in α-synucleinopathy including propagation. There have been reported genetic link between *GBA1* and other α-synuclein diseases such as dementia with Lewy body or multiple system atrophy (MSA). Our data provides evidence that GCase supplementation can rescue α-synucleinopathy in disease models and is therefore a promising approach for the treatment of not only PD, but more general α-synucleinopathies [2, 34].

## Conclusions

In summary, using *in vitro* and *in vivo* models, we have demonstrated that GCase supplementation can restore GD phenotypes including lipid imbalance and motor dysfunction and plays a direct role in the clearance of α-synuclein pathology. We have compared AAV5 and AAV9 highlighting the difference in liver distribution after i.c.v. administration. The two ubiquitous promoters CBh and EFS performed similar in most assays, with the difference of EFS performing better in iPSC neuronal cells and *in vivo* for restoring motor dysfunction in the 4L/PS-NA GD mouse model, emphasizing that a detailed analysis of regulatory elements and AAV serotype should be done when developing an AAV gene therapy. GCase supplementation is a promising approach not only for GD and GBA-PD, but also for idiopathic PD and α-synucleinopathies in general.

## Acknowledgements

We would like to thank Dr. Masato Hasegawa in Tokyo Metropolitan Institute of Medical Science for providing mouse pre-formed fibrils (PFFs). We like to thank the team at QPS Austria GmbH, namely David Amschl and Ewald Auer for performing the 4L/PS-NA mouse experiments.

We thank the following members of Takeda Pharmaceutical Company Limited, Ceri Davies, Hideki Matsui, Yusuke Naito, Kentaro Otake, Tomoyuki Kakizume, Masaaki Nishimura, Saku Miyamoto, Jeong-Ho Oak, Masaaki Kakehi, and David Happel for scientific discussions. We would like to thank Ashutosh Gupta, Brian Garner, and Bhanu Dasari for AAV vector production and AAV analytics, and Yurie Sayama, Kotaro Ogawa, and Takashi Obara for technical assistance. We thank Xin Liu for project management.

## Supporting information

**S1 Fig. Brain sampling method using brain slicer in *Gba1* D409V KI study.** (A) Illustration of sampling scheme. (B) Illustration of the injection site (×) and positions where the dissections blades are inserted for tissue isolation. (C) Illustration of cross section of (2). (D) Illustration of cross section of (3). (E) Pictures of the positions of blades on a brain slicer and sampling procedures.

**S2 Fig. *Gba1* D409V KI mouse study treated with AAV5- or AAV9-*GBA1*.** (A) Human *GBA1* mRNA in brain and liver was measured by using qPCR method. Each graph represents the mean ± S.E.M. (n = 4 or 8). The Y-axis shows logarithmic scale. Statistical analyses were performed by Dunnett analysis. *: <0.05; **: <0.01; ***: <0.001, compared to Group 2 and Group 5 for batch 1 study and batch 2 study, respectively. (B) Correlation analysis between VG and mRNA. Both X-axis and Y-axis are logarithmic scale.

**S3 Fig. 4L/PS-NA mouse model treated with AAV5-*GBA1*** (A) Graph represents body weights per group over time. Animals that died prematurely were excluded from the analysis. (B) VG in brain and liver was measured by qPCR method. The Y-axis shows logarithmic scale. (C) Human *GBA1* mRNA in brain and liver was measured by using qPCR method. The Y-axis is logarithmic scale. Each graph represents the mean ± S.E.M. (n = 8–10). (D) Correlation analysis between VG and mRNA in each tissue. Both X-axis and Y-axis show logarithmic scale.

**S4 Fig. Phospho-α-synuclein reduction in mouse primary neuron cells treated with AAV9-*GBA1*.** (A) Representative images of cells treated with AAV9-CBh-*EGFP*: pSer129 α-synuclein aggregates (red), nuclear (blue), and EGFP (green) are shown. (B) Representative images of cells treated with AAV9-CBh-*GBA1* or AAV9-EFS-*GBA1*: pSer129 α-synuclein aggregates (red) and nuclear (blue) are shown.

**S5 Fig. A53T M83 mouse model treated with different dose of CBE.** (A) Table for CBE dose finding. (B) Graph represents body weights per group over time. (C) GCase activity in cortex, hippocampus, and striatum was analyzed and the graph shows mean values per group, and (D) GlcSph levels were determined, see tables S7 and S8 for mean values per group. The Y-axis is logarithmic scale.

**S6 Fig. CBE-treated A53T M83 mouse model with AAV5-*GBA1* administration.** (A) Graph represents body weights per group over time. (B) Western (WES) images of monomeric (∼19 kDa) and HMW α-synuclein (48–230 kDa) in Triton X-100-insoluble fraction of striatum from groups is shown. Each lane represents each mouse striatum sample data. (C) Graph represents AUC of monomeric α-synuclein normalized with the average of Group 1 by WES. Statistical analyses were performed by Dunnett analysis. *: <0.05; **: <0.01; ***: <0.001 compared to group 2 (sham, CBE).

**S1 Table. Mean GCase Activity and Fold Change in Fig 3B**. S1 Table lists the mean values ± S.E.M. for GCase activity per group and mean fold change for Fig 3B.

**S2 Table. Mean GlcSph Level and Fold Change in Fig 3C**. S2 Table for the mean values ± S.E.M. for GlcSph levels per group and mean fold change for Fig 3C.

**S3 Table. Mean GCase Activity and Fold Change in Fig 4B**. S3 table summarizes the mean value ± S.E.M. for GCase activity per group and mean fold change for Fig 4B.

**S4 Table. Mean GlcSph Level and Fold Change in Fig 4C**. S4 Table for the mean values ± S.E.M. for GlcSph levels per group and mean fold change for Fig 4C.

**S5 Table. Mean GCase Activity and Fold Change in Fig 6B**. S5 table lists the mean values ± S.E.M. for GCase activity for Fig 6B.

**S6 Table. Mean GlcSph Level and Fold Change in Fig 6C**. S6 Table lists the mean values ± S.E.M. for GlcSph levels per group and mean fold change for Fig 6C.

**S7 Table. Mean GCase Activity and Fold Change in S5C Fig.** S7 Table lists the mean values ± S.E.M. for GCase and fold change in S5C Fig.

**S8 Table. Mean GlcSph Level and Fold Change in S5D Fig.** S8 Table list the mean values ± S.E.M. for GlcSph and fold change in S5D Fig.

