## Supplemental Figures (S1-S6) for "AAV delivery of *GBA1* suppresses α-synuclein accumulation in Parkinson’s disease models and restores motor dysfunction in a Gaucher’s disease model"

### S1 Fig.

**A**

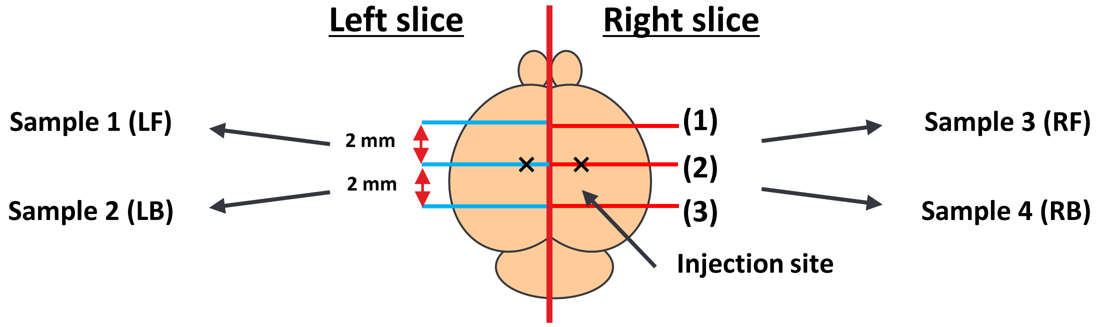


**B**

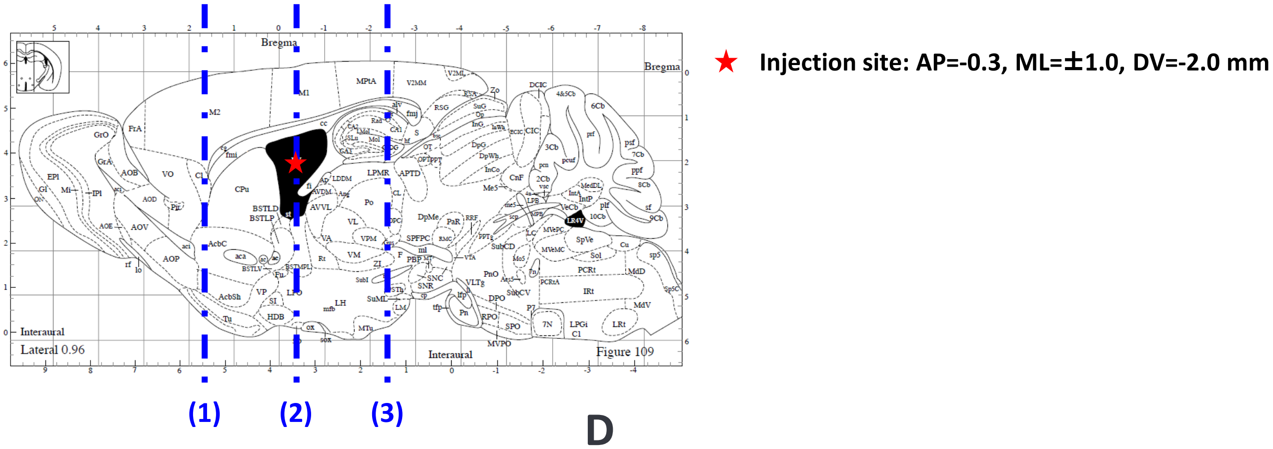


**C**

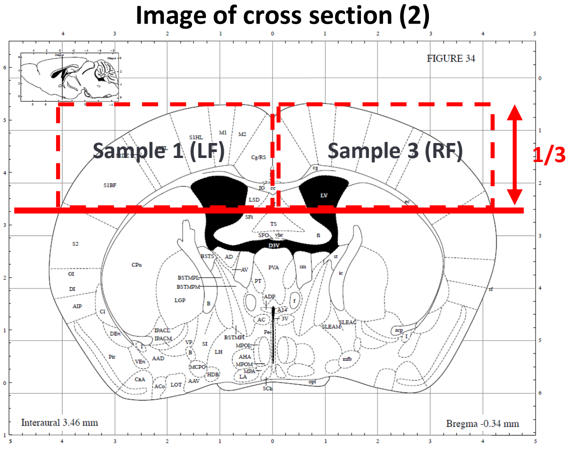


**D**

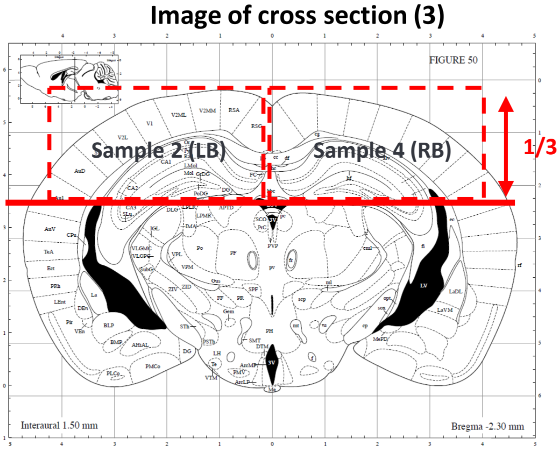


**E**

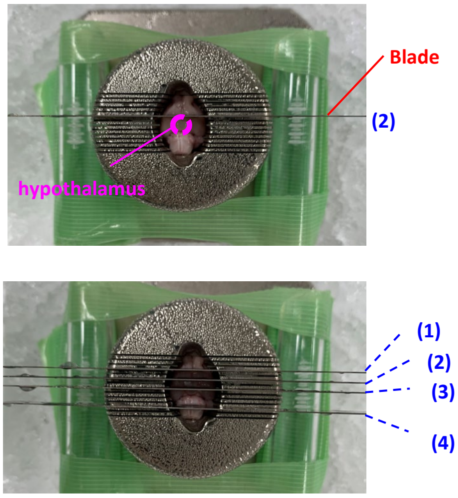


**Step 1**

Place the second blade (position (2)) so that it is just above the tip of the hypothalamus.
(This position is about 0.5 mm before the site of administration, so be careful not to insert it earlier than this.)

**Step 2**

Place the first (position (1)) and third (position (3)) blades 2 mm in front of and behind the second blade (skip one square in the brain matrix).

Placing another blade around the brainstem (position (4)) prevents the brain from shifting when cutting.

**Step 3**

When cutting, use a flat object (e.g. 15-ml tube) to push both ends of the blades evenly and all at once. Cutting one blade at a time can cause the brain to shift.

### S2 Fig.

#### A. hGBA1 mRNA expression

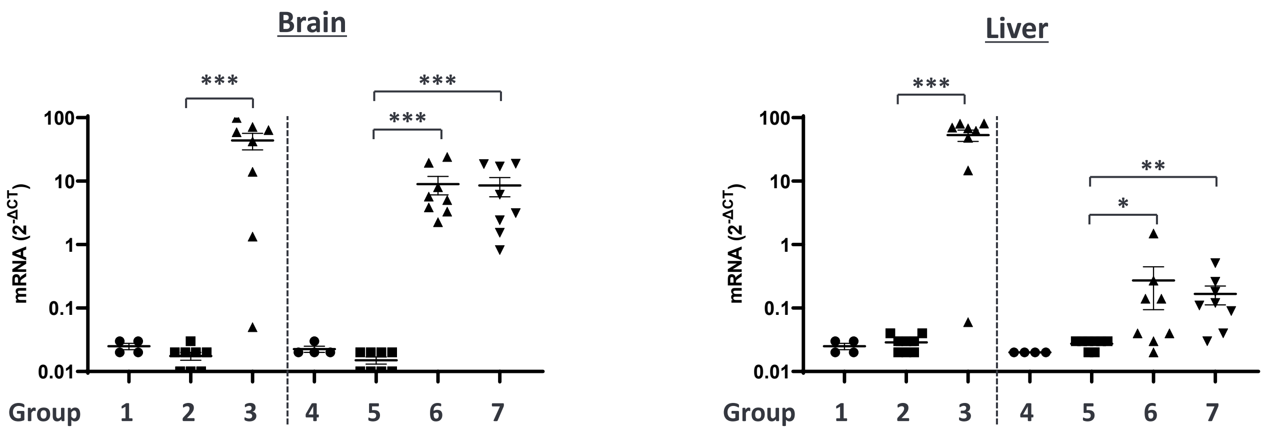


#### B. Correlation between VG and mRNA

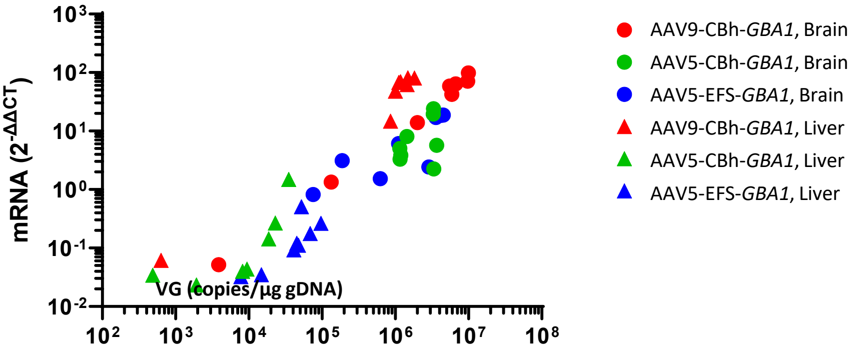

### S3 Fig.

#### A. Body weight over time

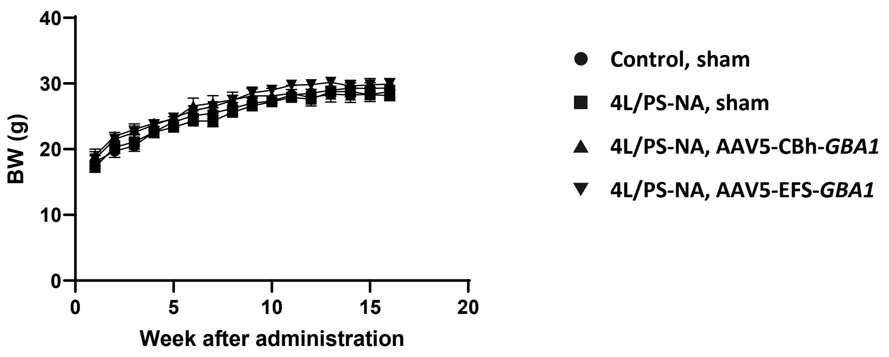


#### B. VG analysis

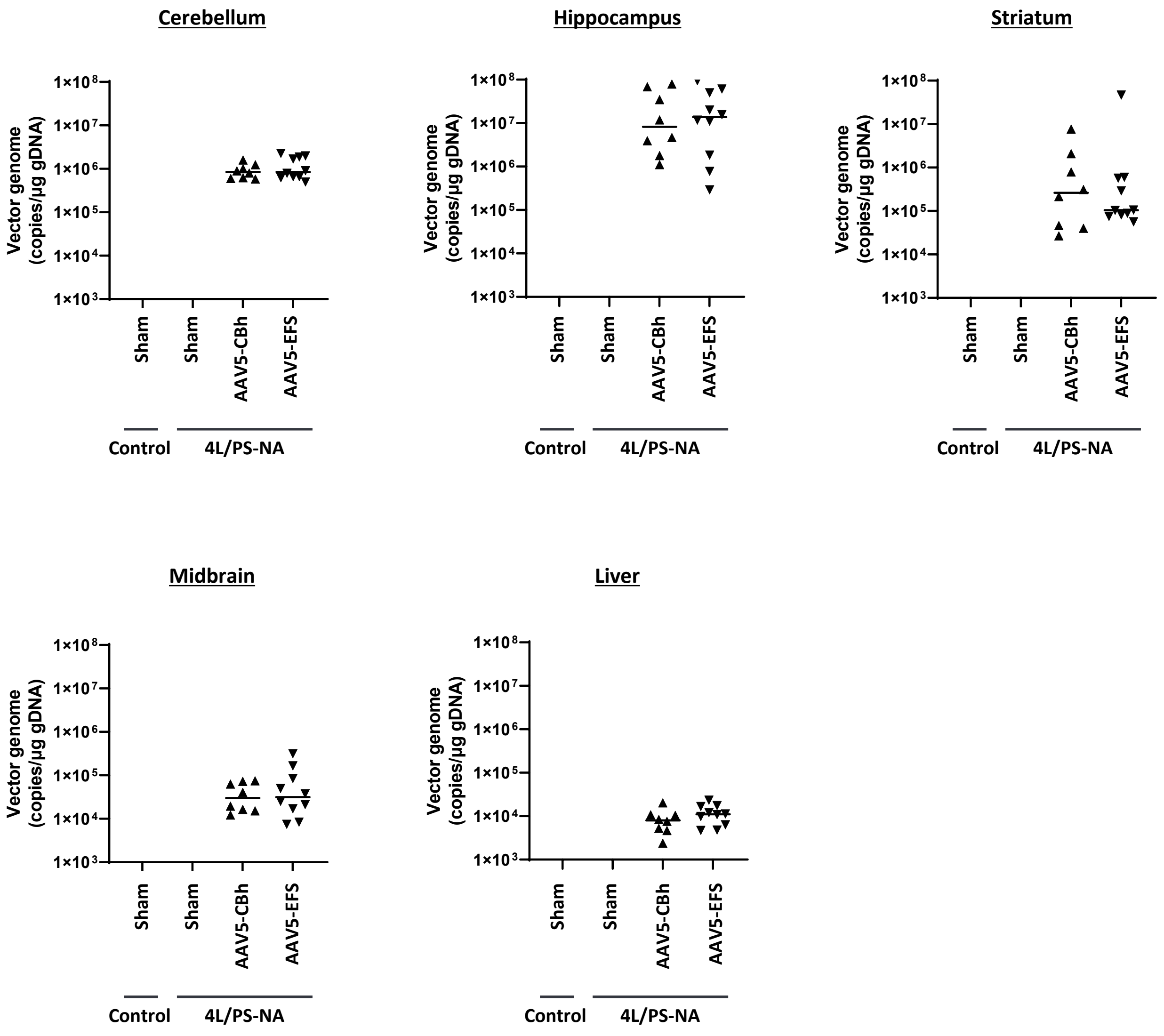

### S3 Fig. (Continued)

#### C. *GBA1* mRNA expression

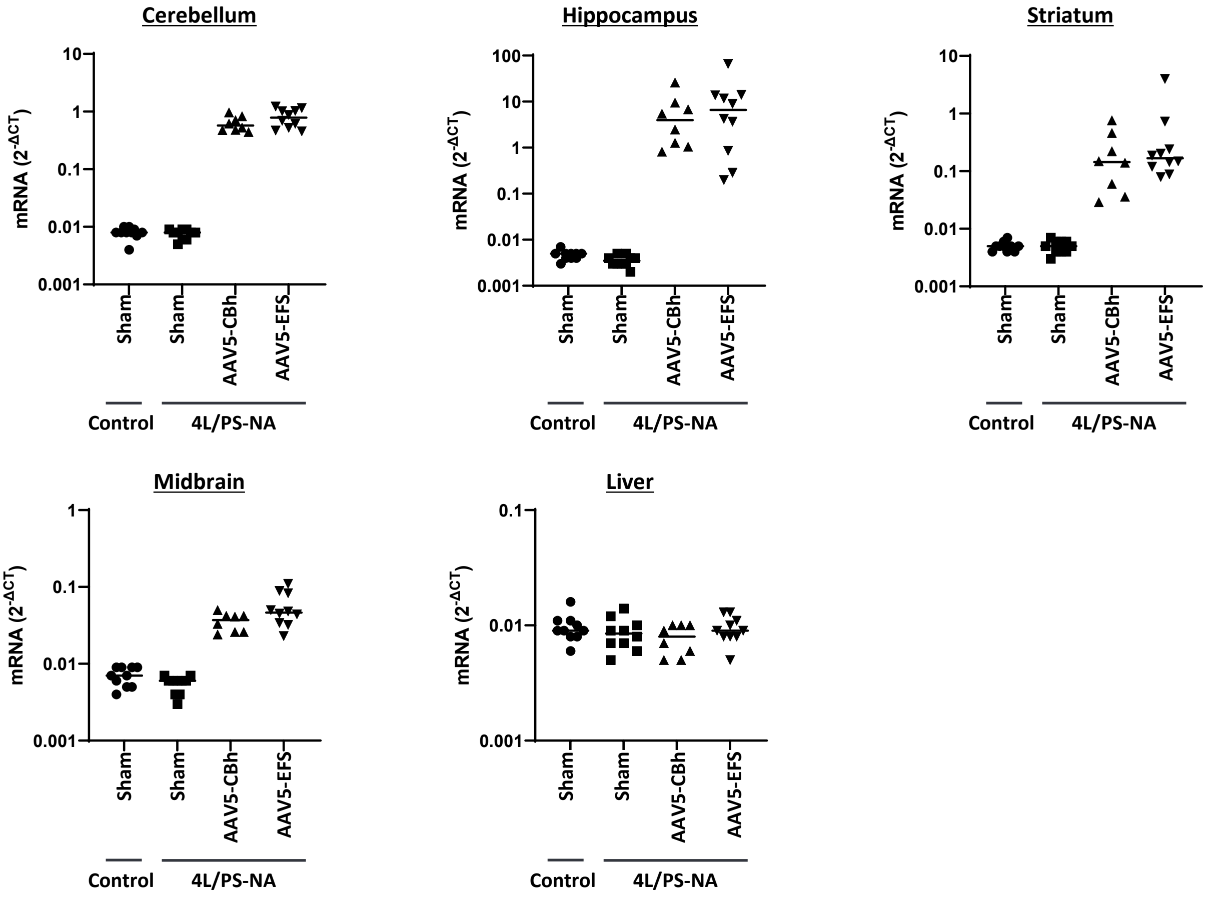


#### D. Correlation between VG and mRNA

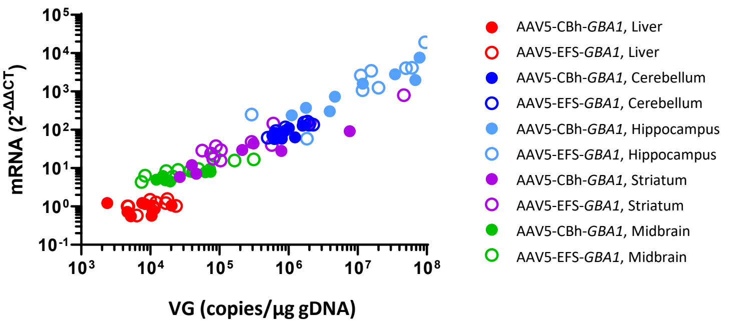

### S4 Fig.

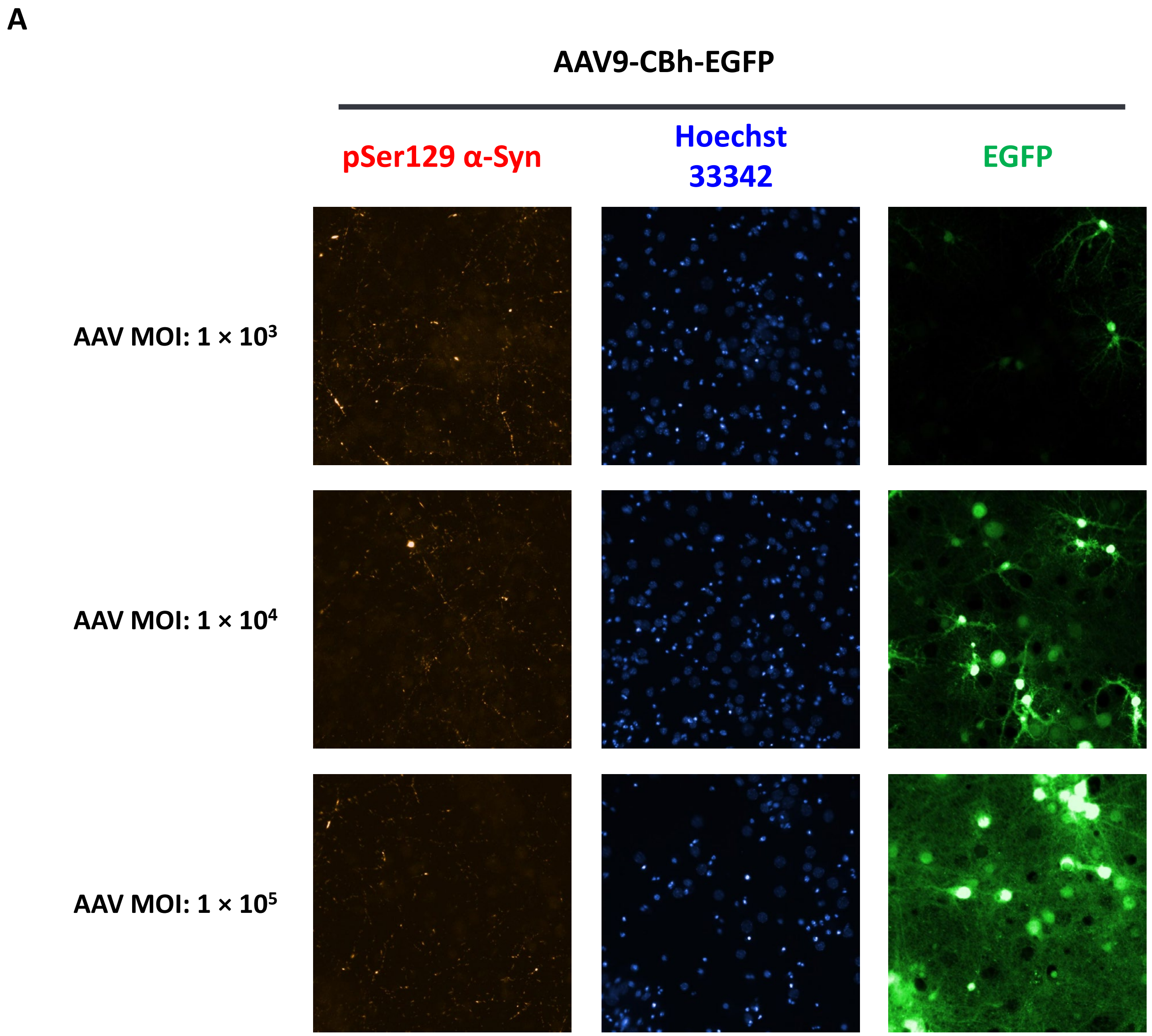
A
AAV9-CBh-EGFP
pSer129 α-Syn
Hoechst 33342
EGFP
AAV MOI: 1 × 10³
AAV MOI: 1 × 10⁴
AAV MOI: 1 × 10⁵

S4 Fig. (Continued)

B

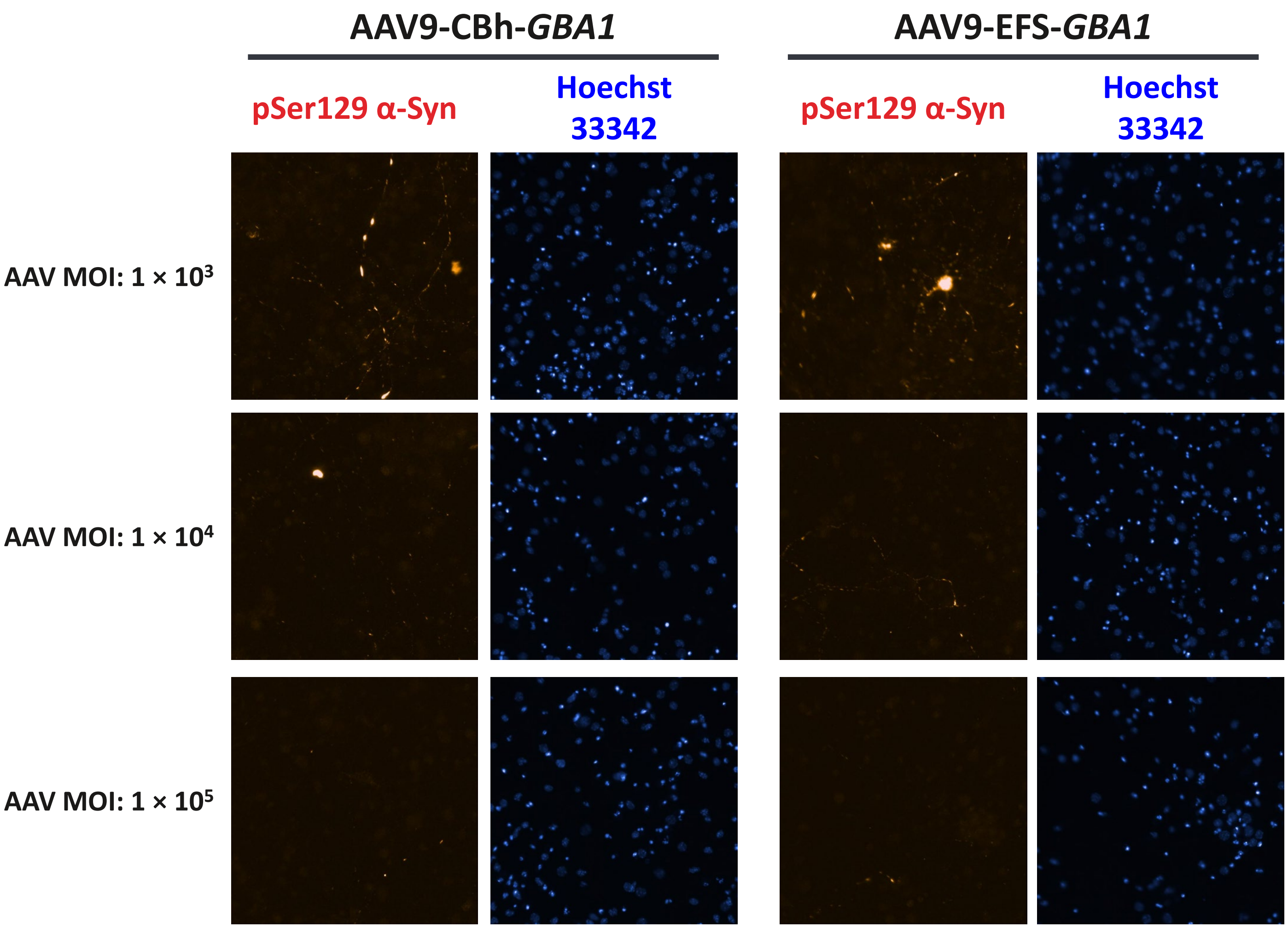

### S5 Fig.

#### A. Treatment Groups

| Group | Animal | CBE treatment | N |
|---|---|---|---|
| 1 | A53T M83 | Saline (i.p., once daily, 10 days) | 6 |
| 2 | A53T M83 | CBE (1 mg/kg, i.p., once daily, 10 days) | 9 |
| 3 | A53T M83 | CBE (5 mg/kg, i.p., once daily, 10 days) | 9 |
| 4 | A53T M83 | CBE (25 mg/kg, i.p., once daily, 10 days) | 9 |

#### B. Body weight over time

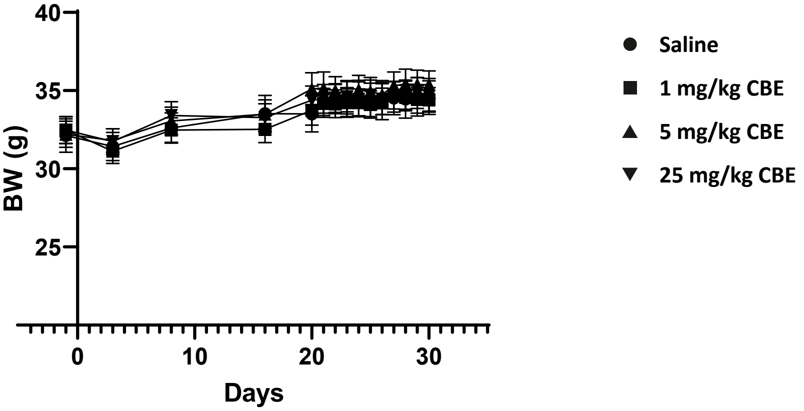


#### C. GCase activity in brain

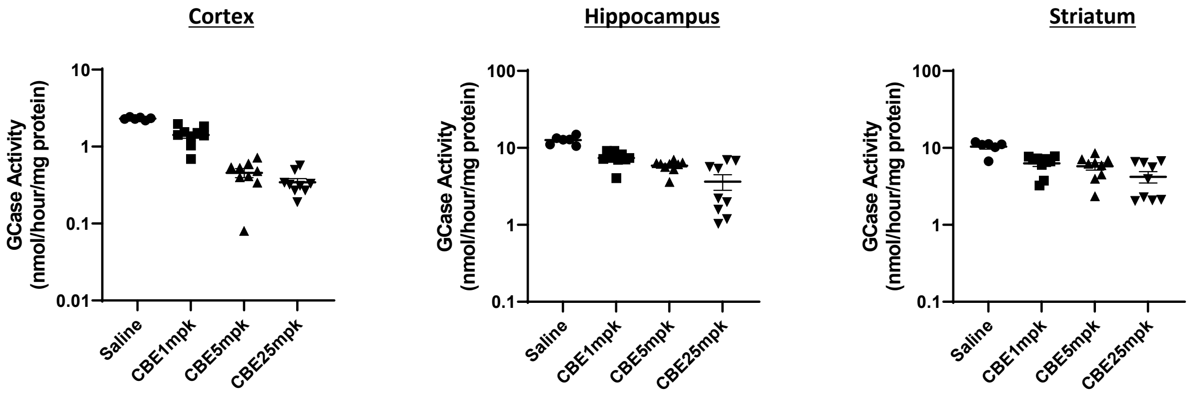


#### D. GlcSph accumulation

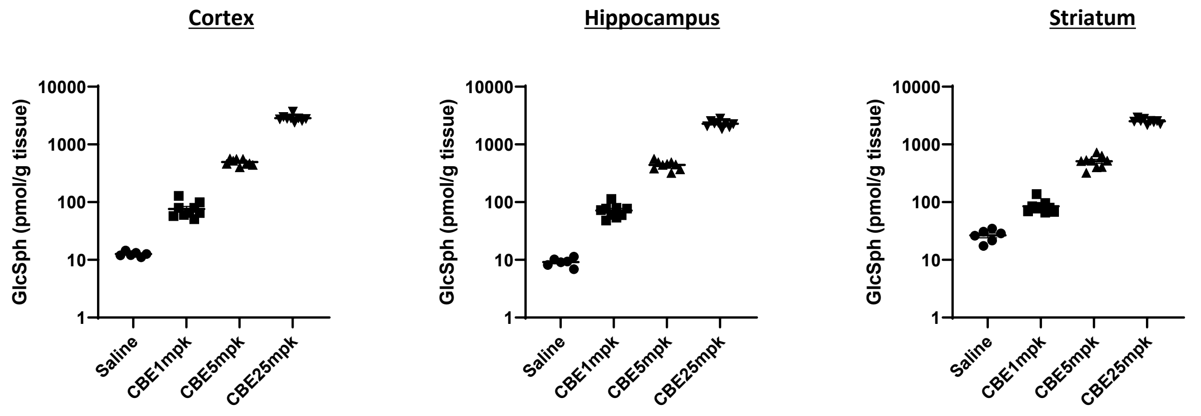

### S6 Fig.

#### A. Body weight over time

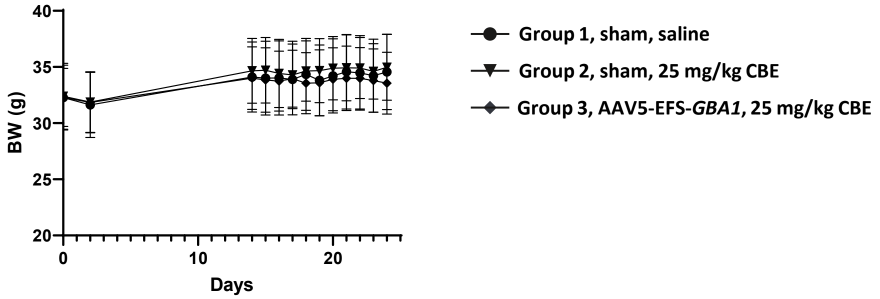


#### B. WES Image (Striatum)

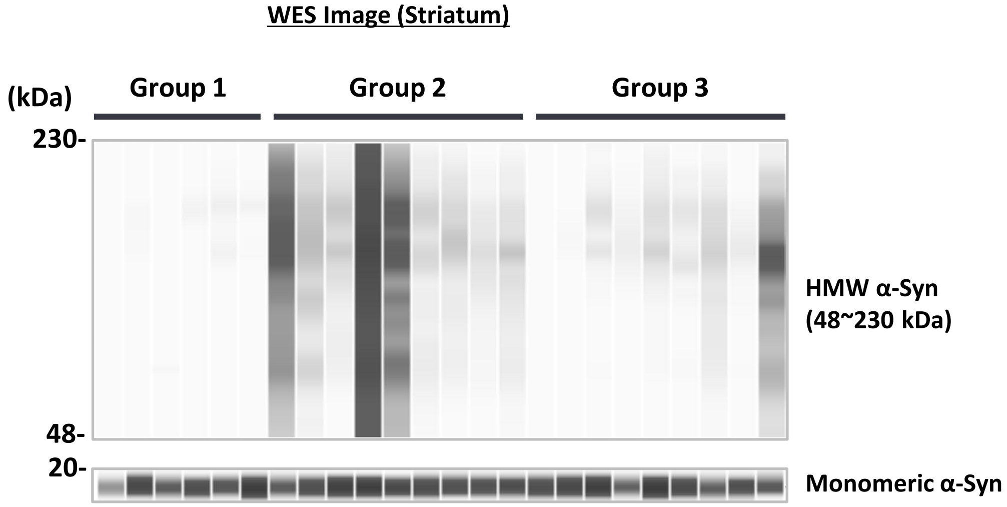


#### C. Monomeric α-Syn quantification (Striatum)

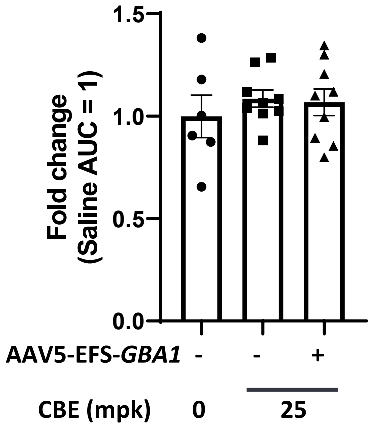
