## Supplemental Tables (S1-S8) for "AAV delivery of *GBA1* suppresses α-synuclein accumulation in Parkinson’s disease models and restores motor dysfunction in a Gaucher’s disease model"

### S1 Table. Mean GCase Activity and Fold Change in Fig 3B.

| Mean GCase Activity ± SEM (nmol/h/mg protein) by AAV9-*GBA1* | | | Mean Fold Increase in GCase activity relative to Group 2 |
|---|---|---|---|
| Group 1 | Group 2 | Group 3 | Group 3 |
| 10.8 ± 0.5 | 5.9 ± 0.2 | 39.8 ± 12.8 | 6.7 |

| Mean GCase Activity ± SEM (nmol/h/mg protein) by AAV5-*GBA1* | | | | Mean Fold Increase in GCase activity relative to Group 5 | |
|---|---|---|---|---|---|
| Group 4 | Group 5 | Group 6 | Group 7 | Group 6 | Group 7 |
| 15.1 ± 0.3 | 9.5 ± 0.2 | 18.5 ± 2.4 | 18.5 ± 3.2 | 1.9 | 1.9 |

### S2 Table. Mean GlcSph Level and Fold Change in Fig 3C.

| | Mean GlcSph level ± SEM (pmol/g tissue) by AAV9-*GBA1* | | | Mean Fold decrease in GlcSph accumulation relative to Group 2 |
|---|---|---|---|---|
| | Group 1 | Group 2 | Group 3 | Group 3 |
| Brain | 7.7 ± 0.7 | 459.1 ± 50.2 | 129.5 ± 57.4 | 3.5 |
| Liver | 105.1 ± 57.4 | 3210.5 ± 75.6 | 794.1 ± 559.8 | 4.0 |

| | Mean GlcSph quantity ± SEM (pmol/g tissue) by AAV5-*GBA1* | | | | Mean Fold decrease in GlcSph accumulation relative to Group 5 | |
|---|---|---|---|---|---|---|
| | Group 4 | Group 5 | Group 6 | Group 7 | Group 6 | Group 7 |
| Brain | 10.4 ± 1.0 | 458.1 ± 42.0 | 108.4 ± 22.8 | 130.5 ± 30.7 | 4.2 | 3.5 |
| Liver | 14.3 ± 8.6 | 3206.3 ± 236.1 | 2410.8 ± 295.7 | 3022.9 ± 189.2 | 1.3 | 1.1 |

### S3 Table. Mean GCase Activity and Fold Change in Fig 4B.

| | Mean GCase Activity ± SEM (nmol/hour/mg protein) | | | | Mean Fold Increase in GCase activity relative to Group 2 | |
|---|---|---|---|---|---|---|
| | Group 1 | Group 2 | Group 3 | Group 4 | Group 3 | Group 4 |
| Cortex | 1.55 ± 0.04 | 2.19 ± 0.07 | 6.80 ± 2.75 | 4.47 ± 0.85 | 3.1 | 2.0 |
| Hippocampus | 0.98 ± 0.03 | 1.23 ± 0.07 | 66.63 ± 24.03 | 78.99 ± 30.07 | 54.0 | 64.1 |
| Striatum | 0.77 ± 0.05 | 1.42 ± 0.08 | 2.76 ± 0.51 | 1.86 ± 0.20 | 1.9 | 1.3 |

### S4 Table. Mean GlcSph Level and Fold Change in Fig 4C.

| | Mean GlcSph quantity ± SEM (pmol/g tissue) or (pmol/mL in CSF) | | | | Mean Fold decrease in GlcSph relative to Group 2 | |
|---|---|---|---|---|---|---|
| | Group 1 | Group 2 | Group 3 | Group 4 | Group 3 | Group 4 |
| Cortex | 1306.7 ± 27.9 | 7210.2 ± 251.0 | 3209.1 ± 342.4 | 3252.7 ± 429.6 | 2.2 | 2.2 |
| Cerebellum | 1317.1 ± 91.1 | 13011.9 ± 606.1 | 7863.5 ± 711.8 | 9675.0 ± 366.3 | 1.7 | 1.3 |
| CSF | 0.090 ± 0.020 | 0.695 ± 0.055 | 0.339 ± 0.067 | 0.427 ± 0.074 | 2.1 | 1.6 |
| Liver | 362.7 ± 15.3 | 5653.1 ± 327.3 | 5940.3 ± 318.6 | 6146.4 ± 224.2 | 1.0 | 0.9 |

### S5 Table. Mean GCase Activity and Fold Change in Fig 6B.

| Mean GCase Activity ± SEM (nmol/h/mg protein) by AAV9-*GBA1* | | | Mean Fold Increase in GCase activity relative to Group 2 |
|---|---|---|---|
| Group 1 | Group 2 | Group 3 | Group 3 |
| 6.7 ± 0.2 | 3.6 ± 0.4 | 10.5 ± 3.4 | 2.9 |

### S6 Table. Mean GlcSph Level and Fold Change in Fig 6C.

| Mean GlcSph quantity ± SEM (pmol/g tissue) | | | Mean Fold decrease in GlcSph relative to Group 2 |
|---|---|---|---|
| Group 1 | Group 2 | Group 3 | Group 3 |
| 29.5 ± 0.7 | 2945.9 ± 257.6 | 1748.0 ± 257.0 | 1.7 |

### S7 Table. Mean GCase Activity and Fold Change in S5C Fig.

| | Mean GCase Activity ± SEM (nmol/hour/mg protein) | | | | Mean Fold decrease in GCase activity relative to Saline group | | |
|---|---|---|---|---|---|---|---|
| | Saline | 1 mg /kg CBE | 5 mg /kg CBE | 25 mg /kg CBE | 1 mg /kg CBE | 5 mg /kg CBE | 25 mg /kg CBE |
| Cortex | 2.32 ± 0.04 | 1.42 ± 0.13 | 0.46 ± 0.06 | 0.34 ± 0.04 | 1.6 | 5.1 | 6.7 |
| Hippocampus | 12.59 ± 0.65 | 7.40 ± 0.51 | 5.86 ± 0.33 | 3.65 ± 0.84 | 1.7 | 2.1 | 3.5 |
| Striatum | 10.38 ± 0.77 | 6.30 ± 0.56 | 5.78 ± 0.62 | 4.21 ± 0.71 | 1.6 | 1.8 | 2.5 |

### S8 Table. Mean GlcSph Level and Fold Change in S5D Fig.

| | Mean GlcSph quantity ± SEM (pmol/g tissue) | | | | Mean Fold increase in GlcSph accumulation relative to Saline Group | | |
|---|---|---|---|---|---|---|---|
| | Saline | 1 mg /kg CBE | 5 mg /kg CBE | 25 mg /kg CBE | 1 mg /kg CBE | 5 mg /kg CBE | 25 mg /kg CBE |
| Cortex | 12.6 ± 0.5 | 76.2 ± 8.2 | 494.1 ± 20.2 | 2845.3 ± 122.8 | 6.0 | 39.2 | 225.9 |
| Hippocampus | 9.2 ± 0.6 | 71.4 ± 6.5 | 441.3 ± 24.6 | 2275.3 ± 96.6 | 7.8 | 48.1 | 248.1 |
| Striatum | 26.6 ± 2.6 | 84.8 ± 7.4 | 509.6 ± 40.9 | 2521.7 ± 80.6 | 3.2 | 19.1 | 94.7 |
